# Inferential planning in the frontal cortex

**DOI:** 10.1101/2025.11.26.690672

**Authors:** Francesco Donnarumma, Thomas Parr, Karl Friston, James Whittington, Giovanni Pezzulo

## Abstract

How the brain plans and maintains sequences of future actions remains a central question in systems neuroscience. Recent studies in the frontal cortex have revealed that multiple elements of a sequence are represented simultaneously in separable neural subspaces, challenging classical serial models of sequential planning. Here, we show that these representations emerge naturally under inferential planning in which sequential actions are inferred from sensory evidence and goals. Using a hierarchical generative model, we reproduce key neural phenomena observed in primate frontal cortex, including the simultaneous activation of multiple plan elements, the emergence of (almost) orthogonal ‘memory’ subspaces, and their reuse across forward and backward sequence tasks. Our approach provides a mechanistic account of how probabilistic inference over control states gives rise to distributed and dynamic neural representations of plans. This framework not only unifies previously disparate findings on planning, working memory, and motor preparation, but also generates novel, testable predictions about the dynamics of active inference, the role of sensory subspaces, and the impact of uncertainty on sequence processing.

## 1 Introduction

The ability to plan is central to intelligent behaviour. Planning involves sequencing a series of actions that can later be enacted in the world. While we are beginning to understand sequence processing in the brain mechanistically (Mattar & Lengyel, 2022; Miller et al., 2017; Pezzulo et al., 2019; Balaguer et al., 2016), our understanding of how plans are formed, and their relationship to sequence processing more broadly, remains incomplete.

The dominant view of planning states that elements of the plan are sequentially sampled one after another, i.e., the plan is generated in series with the neural population representing just one action at any time-point. This view is consistent with many existing models for sequence understanding, as well as planning, such as recurrent neural networks (Elman, 1990; Botvinick & Plaut, 2006; Maass et al., 2002; Ganguli et al., 2008; Jensen et al., 2024; Di Antonio et al., 2024), probabilistic generative models (Parr et al., 2024; Pezzulo et al., 2017, 2014; George et al., 2021), or heteroclinic channels (Rabinovich et al., 2014). Representing plans and sequences sequentially has been able to account for a variety of different neural representations, from motor sequences as dynamical systems (Mante et al., 2013; Sussillo et al., 2015) to hippocampal replay sampled from a generative model (cognitive map) of the environment (Schwartenbeck et al., 2023; Stoianov et al., 2022; Foster, 2017) and the mental planning of multiple tasks reported in the orbitofrontal cortex (Wilson et al., 2014; Schuck et al., 2016; Van de Maele et al., 2024). Further, such formalisms have been combined with reinforcement learning (Stachenfeld et al., 2017; Behrens et al., 2018; Baram et al., 2021), as well as the probabilistic planning literature (George et al., 2021; Raju et al., 2024; Stoianov et al., 2018), to understand goal directed behaviour with models predicting a variety of neural phenomena of the hippocampal formation. Despite their differences, all these models generate sequential transitions between elements—whether spatial positions, memories, or actions. At the neural level, this implies that at any specific point in time, the brain represents only one element of the sequence in ongoing neural activity.

Recently, however, a different type of representation has been uncovered in prefrontal cortex. Here, all elements of a sequence are represented simultaneously in neural activity at any given time, from path planning tasks (Mushiake et al., 2006; Saito et al., 2005; El-Gaby et al., 2024) to sequence memory tasks (Xie et al., 2022; Panichello et al., 2024; Chen et al., 2024) to drawing (Averbeck et al., 2002) and visual working memory tasks (Liu et al., 2024). Each element of the sequence is represented, and ordered, in a distinct set of neural subspaces—termed ‘activation slots’ (Whittington et al., 2025). This allows the entire sequence to be represented simultaneously across the different slots. While this is a different view of sequence representation, it has been shown to have a mathematical equivalence to the sequential view described earlier (Whittington et al., 2025). This raises the tantalising possibility that this new type of representations—slots—can also serve as a neural substrate of planning more broadly.

In this paper, we formalise planning as inference over these ordered subspaces. We show that messages are passed from slot to slot in order to infer the correct state and action held at each time-step of the plan. Using this theory we able to explain a variety of neural findings from prefrontal cortex while animals plan: 1) single neurons coding for specific cursor movements at specific time-steps in the plan (Mushiake et al., 2006); 2) neural subspaces holding arbitrary elements in sequence memory task (Xie et al., 2022); 3) showing that the same subspaces are reused in both forward and backward planning tasks (Chen et al., 2024); 4) inferring plans when the number of sequence elements in a plan differs (Chen et al., 2024). Our theory shows planning and sequence memory to be two sides of the same coin and offers a mechanistic understanding of how an entire plan can be encoded—in neural activity—simultaneously in prefrontal cortex.

## 2 Results

We first introduce an inferential planning model (Section 2.1) used subsequently to reproduce the neural dynamics observed empirically in three studies (Mushiake et al., 2006; Xie et al., 2022; Chen et al., 2024).

### 2.1 A prefrontal model to infer arbitrary plans

We start by considering what it would mean to represent an arbitrary plan simultaneously in a set of neurons (or neuronal populations), and then construct a model that is able to infer such a plan representation.

A plan is a proposed sequence of actions (*a*_*t*_) and states (*s*_*t*_): { (*s*_*t*_, *a*_*t*_)}. Actions, however, can be abstract or higher-level subgoals (e.g., go to A) that are subsequently enacted by motor controls (e.g., move arm left). To account for this, we use a hierarchical model, in which the plan is represented in variables abstracted from motor commands (in Level 2), while the motor commands themselves are represented in Level 1 (Figure 1A).

**Figure 1.**
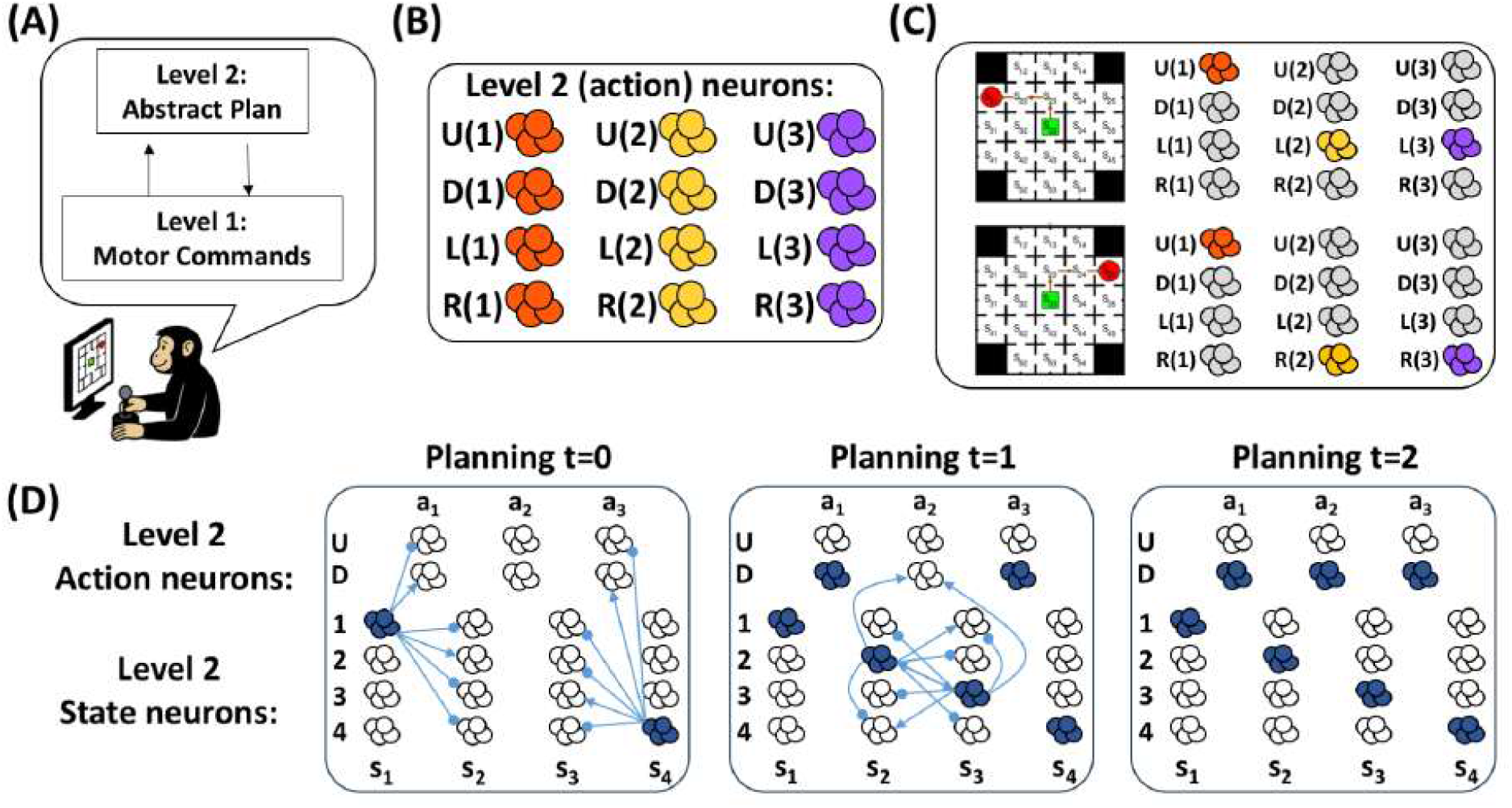
Schematic illustration of the hierarchical inferential planning model. (A) The model comprises two interconnected levels. Level 2 (planning level) encodes beliefs about plans—abstract sequences of actions (and states) over time. Level 1 (sensorimotor level) encodes beliefs about concrete motor commands (and sensory states), which generate observable outcomes in the environment. Arrows indicate the bidirectional exchange of information: top–down predictions from Level 2 guide action selection at Level 1, while bottom–up sensory evidence updates beliefs at both levels. This architecture supports both forward and backward inference over time, allowing the system to infer, maintain, and flexibly revise entire action sequences. (B) Within Level 2, neural populations encode beliefs about abstract actions Up, Down, Left, and Right joystick movements (and corresponding states, omitted for clarity) at multiple time points; for example, U1 is Up at time point 1, U2 is Up at time point 2, etc. These are maintained and updated in parallel through reciprocal message passing (Section 4). (C) Example of Level 2 neural representations of two plans: {*Up, Left, Left*} and {*Up, Right, Right}* . (D) Intuitive understanding of the message passing process in which plans are inferred. In this schematic, we show both example state neurons and action neurons of Level 2 (for a simplified task compared to the other panels). First, a start and goal state are presented and inferred by populations of state neurons (blue) in the left box. Then, through multiple iterations (boxes from left to right), the full plan is inferred by action neurons, while state neurons simultaneously represent expected states along the trajectory from start to goal. The inference of action and state nodes is constrained by prior knowledge about start and goal states, as well as the consequences of actions (i.e., the transition function or connectivity among neural populations), and is implemented via local message passing between neural populations, here schematically represented by blue arrows with excitatory (arrowheads) and inhibitory (circles) roles. For example, inferring a specific goal state increases (excites) the probability of action nodes that achieve it, while decreasing (inhibiting) the probability of action nodes that diverge from it. Similarly, inferring a specific action increases the probability of state nodes that result from taking that action. Ultimately, the sequence of action and state neurons that is inferred is the most likely path from start to goal. Please note that this figure presents a simplified model intended to build intuition, omitting the mathematical details of how the most likely sequence of actions and states is inferred. See Section 4 for the full formal implementation based on active inference (Parr et al., 2022).

For Level 2 neurons to represent arbitrary full plans (complete sequences of action and states), multiple copies of action and state neurons are required in which each copy corresponds to potential actions and states at each time-point in the plan (Figure 1B). Correspondingly, this means the whole plan is simultaneously encoded by neural activity. For example, if each planning problem involves a sequence of three joystick movements—example sequences being {*Up, Left, Left*} or {*Up, Right, Right*} (with corresponding states at each time-point)—there must be three pools of neurons each able to represent any state and any action at their corresponding time-point (Figure 1C).

This simultaneous representation of all elements in the plan allows for *arbitrary* plans to be represented. This is fundamentally different to formalisms where elements of a plan are represented sequentially, one after another, in the same neuron or population. While that setup can represent *specific* sequences using neural dynamics (provided the appropriate synaptic connections), it does not have the flexibility to represent *arbitrary* plan sequences. Furthermore, the encoding of the first action cannot be revised or contextualised by the last action, because the first action is encoded in the past, when the last action is represented.

The ensuing simultaneous sequence element setup bears a remarkable resemblance to the ‘activation slots’ found in prefrontal cortex (Whittington et al., 2025). However, current models of ‘activation slots’ are limited to understanding just sequential memory tasks. In this paper, we show that the same computational architecture can solve arbitrary planning tasks. In order to form a plan, however, inference is required. This may involve inferring the path between a start and goal state, though more generally it is the process of combining observations with task statistics to infer optimal behaviour (we provide examples of this later in the paper). Regardless of the setting, we employ variational Bayes for planning as inference (Attias, 2003; Botvinick & Toussaint, 2012; Friston et al., 2017a; Parr et al., 2022). Thus the higher level (Level 2) encodes beliefs about plans or *policies*, while the lower level (Level 1) encodes beliefs about concrete sensorimotor actions that implement these plans—by moving the agent from one location to the next—interacting directly with the environment.

Information flows bidirectionally between the two levels and with the environment. Predictions about planned sequences flow downward from Level 2 to Level 1, guiding the selection of sensorimotor actions that are executed in the environment. In turn, sensory feedback and action outcomes generate bottom-up signals that update beliefs at Levels 1 and then 2. This bidirectional exchange allows the system to continuously align planned sequences with sensory evidence, ensuring coherent behaviour across hierarchical levels of control. Within each level, neural populations represent probabilistic beliefs about hidden states, and their interactions implement probabilistic belief updating via a message passing method, called belief propagation (Pearl, 2022; George & Hawkins, 2009; George et al., 2025; Friston et al., 2017a,b; Parr et al., 2019a).

In sum, we consider a (Level 2) neural population which consists of copies (subspaces) of state and action representations, each able to represent an arbitrary state and action. To infer what action and state should go in each subspace (i.e., infer the plan), we use a neuronally plausible message passing algorithm (see Figure 1D for an intuitive illustration). Critically, the simultaneous representation of all time-steps in the plan is essential to support the reciprocal message passing—i.e.,belief propagation—that implements this sequential planning. See Section 4 for a formal description of the model and implicit message passing.

We now show a series of results (Sections 2.2, 2.3, 2.4) in which our model recapitulates the prefrontal neural activity of several experimental studies. In each case, we show neural population dynamics corresponding to the inference process of Level 2 abstract plans. Where specified, we also show dynamics corresponding to inference of Level 1 specific actions.

### 2.2 Planning sequences of cursor movements

First we consider a planning task (Mushiake et al., 2006) in which monkeys were trained to control a cursor to navigate a two-dimensional maze, moving from a central position to a cued goal location (Figure 2A-B). Their key finding was that, during the preparatory period preceding movement, simultaneous representations of multiple future cursor movements were recorded in distinct neuronal populations within the monkey lateral prefrontal cortex, each selective for a specific movement at a specific time step, e.g., the Right cursor movement at the 1st time step, the Left cursor movement at the 2nd time step, or the Left cursor movement at the 3rd time step (Figure 2C). In contrast, neural activity in the primary motor cortex primarily reflected actual arm movements.

**Figure 2.**
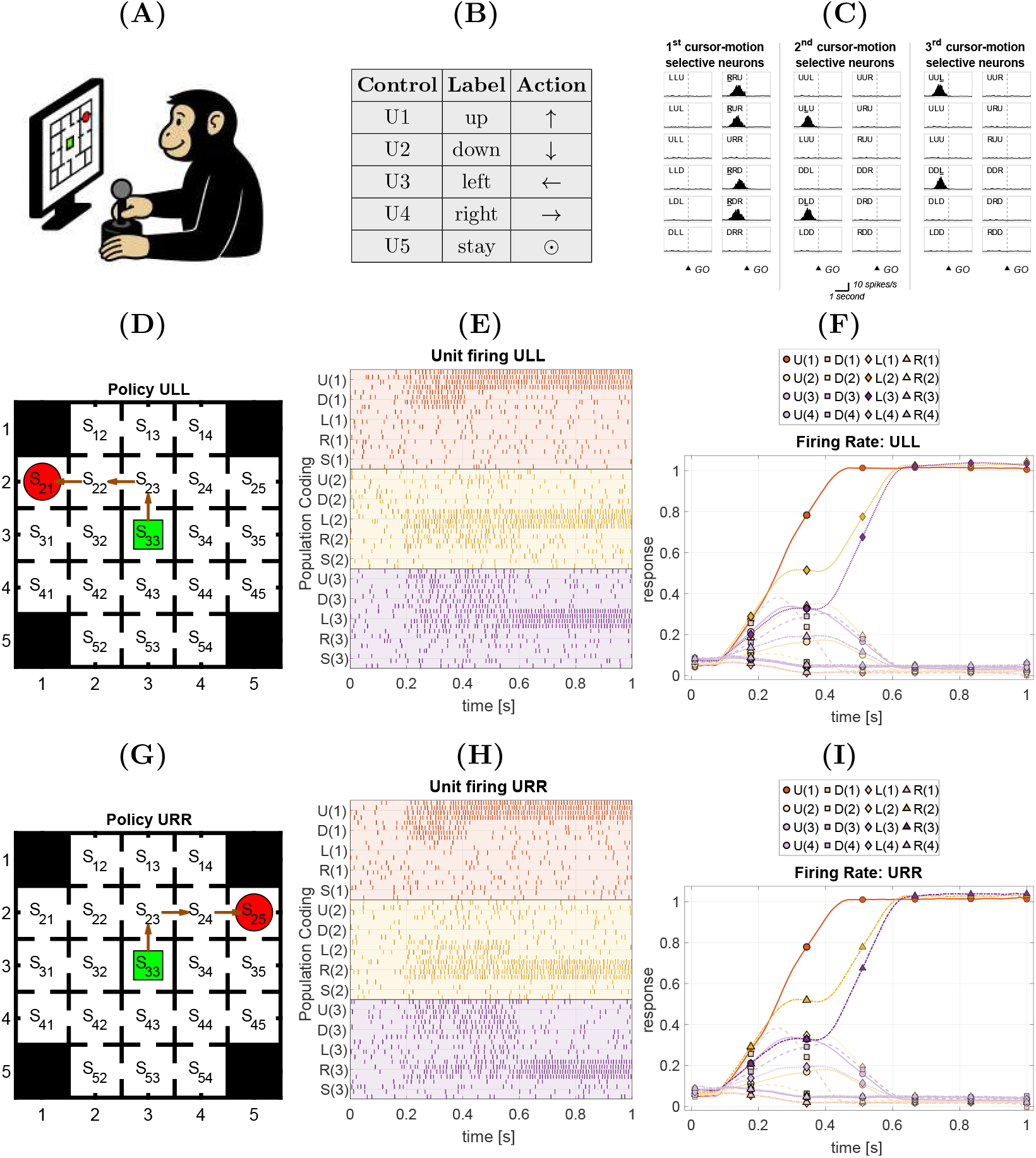
Planning a sequence of three cursor movements in a maze. This simulation replicates the experimental setup of (Mushiake et al., 2006). (A-B) Monkeys navigated a 5 × 5 ‘maze’ by controlling a joystick. They could perform five possible cursor actions: Up, Down, Left, Right, and Stay. (C) Schematic illustration of the results reported in (Mushiake et al., 2006, Figure 3), showing three neurons selective for the Right cursor movement at 1st time step, the Left cursor movement at the 2nd time step, and the Left cursor movement at the 3rd time step. These neurons correspond to R(1), L(2) and L(3) in our simulations. (D-F) and (G-I): two example maze navigation problems, consisting in moving from the start (green square) to the goal state (red circle). Each white cell represents a valid navigable state, whereas black cells indicate blocked states. Walls are depicted as solid lines, and open lines (*doors*) represent allowed transitions. For each configuration, the successful strategy (shortest policy) is highlighted with arrows. (D) In this example maze, the planned policy is Up-Left-Left (*ULL*). (E) Raster plots showing simulated neural activity, with four neurons coding for each of the 5 control states (*Up, Down, Left, Right, Stay*) at Level 2, at each each of the three time steps. Neurons coding for actions at the 3 time steps are color coded in red, yellow and violet, respectively. At the end of the simulation, corresponding to the delay period before execution, the policy *ULL* is coded by the simultaneous activation of three populations of neurons, *U* (1), *L*(2) and *L*(3). (F) The same results, but plotted as simulated firing rates of neurons associated with different actions. Neurons representing *U, D, L* and *R* are coded with circles, squares, diamonds, and triangles, respectively. The stay action is not shown for simplicity. (G) In this example maze, the inferred policy is Up-Right-Right (URR). (H-I) Simulated raster plots and firing rates for the policy URR, following the same format as Panels C-D. See the main text for further explanation.

We simulated this task under our hierarchical model (Figure 1). The lower level (Level 1) controls cursor movements, whereas the upper level (Level 2) plans the sequence of cursor movements. Level 2 comprised 25 hidden states representing maze locations (from *S*_11_ to *S*_55_) and 5 control states corresponding to possible cursor movements: Up, Down, Left, Right, and Stay. The simulated neural activity reported below corresponds to the inference of these 5 control states across the 3 time steps of the plan, which we compare with the frontal cortical activity reported by (Mushiake et al., 2006).

We denote time-steps within the plan (but all still represented simultaneously) with parentheses; for instance, the ‘Up’ action at the first, second, and third time steps is represented as U(1), U(2), and U(3), respectively. We use 4 neurons to represent each hidden state, and thus 60 neurons (4 ×5× 3) to represent the 5 possible cursor movements across 3 time steps

Figure 2D illustrates an example problem in which the agent plans a sequence of three cursor movements to navigate from the start location (green; *S*_33_) to the goal location (red; *S*_21_). Figure 2E shows a simulated raster plot representing the inferential process for the five possible cursor movements (Up, Down, Left, Right, Stay) across the three time steps. The 15 horizontal lines represent the simulated firing of 15 neural populations (each comprising four neurons) encoding the five cursor movements at three time steps. For instance, U(1) (first line), U(2) (sixth line), and U(3) (eleventh line) correspond to planning the ‘Up’ movement at the first, second, and third time steps, respectively. Cursor movements at different time steps are color-coded: red, yellow, and violet correspond to the first, second, and third time steps, respectively.

At the start of the planning process (time 0), the agent maintains a relatively flat belief distribution over the first, second, and third planned cursor movements. Upon presentation of the goal location *S*_21_, it updates its beliefs about all three movements. During the early stage of planning (approximately 0.2–0.6), parallel competition occurs among the two available cursor movements at time step 1 (U(1) and D(1)), the two most likely movements at time step 2 (L(2) and R(2)), and the three most likely movements at time step 3 (U(3), L(3), and R(3)). By around 0.6, planning is completed, and the three selected cursor movements (U(1), L(2), and L(3)) exhibit simultaneous, sustained activation, which can serve as a prospective working memory for the plan. However, as time passes—and the plan is executed at the lower level—mnemonic representations of early elements become postdictive.

The dynamics of the inferential process can also be observed in Figure 2F, which shows the average firing rates of the 12 neural populations involved in cursor movement planning (with ‘Stay’ actions omitted). The figure illustrates that the three selected cursor movements (U(1), L(2), and L(3)) gradually increase their activation over time, with the first planned movement (U(1)) rising fastest, reflecting faster convergence during inference. These dynamics resemble a competitive queuing mechanism, in which the first element of a sequence is activated first, but here this pattern emerges spontaneously during inference. It is also apparent that most of the non-selected cursor actions increase slightly before decreasing, as they lose the competition with alternative plans. Unfeasible actions (e.g., L(1) and R(1)) instead show an immediate decrease in activity.

Figures 2G–I illustrate another example problem, in which the agent plans a sequence of three cursor movements (U(1), R(2), and R(3)) from the start location (green; *S*_33_) to the goal location (red; *S*_25_), along with the corresponding inferential dynamics, using the same format as Figures 2D–F. See also the Supplementary Materials for additional illustrations of the neural dynamics associated with inference of hidden states –actions (Supplementary Figure S1) and locations (Supplementary Figure S2) at both Levels 1 and 2.

In summary, these simulations demonstrate how inferential planning dynamics naturally reproduce the simultaneous activation and updating of neural populations encoding cursor movements at different time steps, as observed in the monkey lateral prefrontal cortex (Mushiake et al., 2006).

### 2.3 Inferring and holding in memory a sequence of saccades

Second, we consider a task (Xie et al., 2022) in which monkeys were first shown a sequence of three targets (from six possible targets arranged in a hexagon) and then, after a delay, were required to saccade to the same three targets in order (Figure 3A-D). Unlike the previous task, this requires inferring a plan from observations and maintaining it in working memory. A key finding of this study was that, during the delay period, the prefrontal representation decomposed into three distinct and near-orthogonal subspaces for each element (referred to as ‘ranks’) in the sequence (Figure 3C).

**Figure 3.**
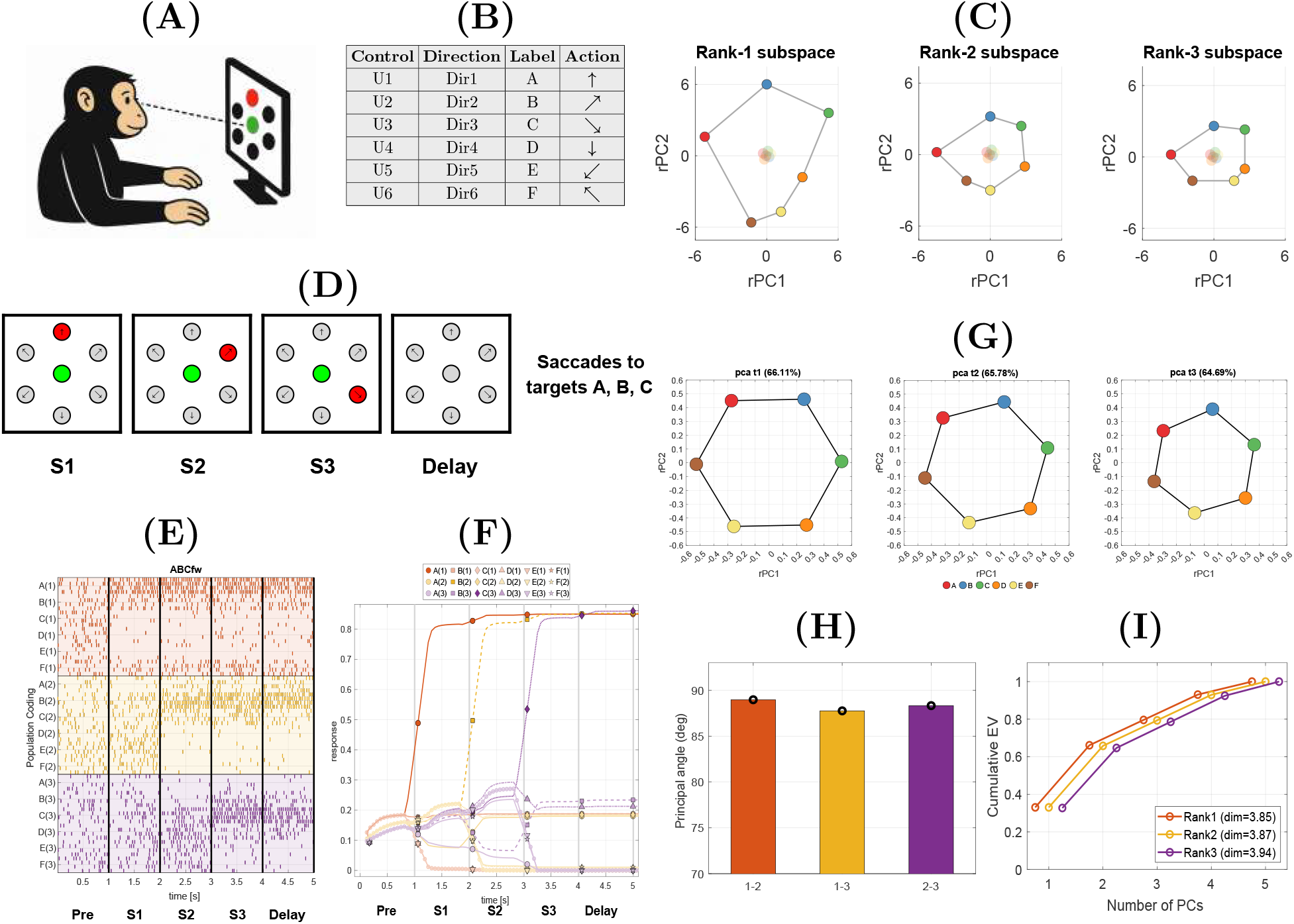
Inferring and holding in memory a sequence of three saccades. This simulation reproduces the experimental setup of (Xie et al., 2022). (A–B) Monkeys were instructed to perform sequences of three saccades toward six possible targets, denoted as {*A*–*F*} . (C) Schematic illustration of the results reported in (Xie et al., 2022, Figure 2). The figure shows the disentangled neural state space representation projected onto three near-orthogonal 2D, step-specific subspaces, one per panel. Within each panel, Black-connected points indicate samples at the end of the delay arranged in a hexagonal geometry; central points correspond to the beginning of the delay. (D) Example of a trial, in which monkeys maintained a central fixation (green square) while they were shown three consecutive targets, *A, B*, and *C*, that they had to fixate in the same order after a delay period. (E-F) Simulated firing rates and spike trains for an example trial, requiring generate sequential saccades to the *A, B*, and *C* targets. (G) Simulated population responses for each combination of step and target during the delay period, projected onto three 2D step-specific subspaces, one per panel; note that these subspaces correspond to the rank-subspaces in (Xie et al., 2022), shown in Panel C. The bottom-left plot shows that the three subspaces are oriented in a near-orthogonal manner in neural state space, as indicated by the large principal angles between them. The bottom-right plot shows the cumulative explained variance along the different steps in neural state space, reflecting the symmetric partitioning of information across the three subspaces. See the main text for further explanation.

Figure 3E–I shows the results of a simulation of the (Xie et al., 2022) experiment. The generative model’s structure is the same as in the previous simulations, but the states at each level differ. Level 1 transiently encodes incoming stimuli and governs eye movements, and is not the focus of our analysis. Level 2 comprises 6 targets for each of the 3 time steps, corresponding to 18 neural populations in Figure 3E. Targets are labelled *A* −*F*, and the number in parentheses denotes the time step. The Level 2 states show two types of mixed selectivity (Rigotti et al., 2013). First, the coding of targets is different across time steps (or “ranks”), which means that, for example, the firing of A(1) reflects a combination of target A and rank 1. This is a special case of non-linear mixed selectivity – a duplication – reflecting the finding in (Xie et al., 2022, Figure S8) that many neurons only participate in one “rank subspace”. Second, while neural populations are strongly tuned to a preferred direction (e.g., A for A(1)), they are also weakly tuned to the neighbour dimensions (B and F). This amounts to linear mixed selectivity – also called cosine directional tuning (Georgopoulos et al., 1982), see Section 4.6 for details.

At the beginning of an example trial (‘Pre’ period), the agent maintains a relatively flat belief distribution over the next targets. During the observation periods ‘S1’, ‘S2’, and ‘S3’ the agent observes targets *A, B*, and *C*, respectively, and incrementally infers that the correct plan is *ABC*. Notably, during ‘S1’ it infers *A*(1) with high probability while simultaneously decreasing the probabilities of *A*(2) and *A*(3), since targets cannot be repeated. During ‘S2’ it maintains its belief about *A*(1) and additionally infers *B*(2). Finally, at ‘S3’ it infers the complete *ABC* plan and maintains it throughout the delay period. The corresponding average firing rates for this inferential process are shown in Figure 3F, in the same format as the first simulation.

To test whether the neural coding in our simulation reproduces the near-orthogonal rank subspaces reported by (Xie et al., 2022), we applied the same analysis approach as the original empirical study, projecting population activity into three 2D step-specific ‘rank-subspaces’. Figure 3G shows the simulated population responses for each target and time step projected onto these subspaces, with the six target locations colour-coded. The results match very well the empirical data. The hexagonal shape of the subspaces is the same as the empirical data (Figure 3C). Furthermore, the large principal angles between the three subspaces (Figure 3H), together with the cumulative explained variance across steps in neural state space (Figure 3I), recapitulate the empirical findings of (Xie et al., 2022) and indicate that the subspaces are nearly orthogonal. Note that variance is concentrated in the leading components of the neural state space (Figure 3I), consistent with (Xie et al., 2022, Figure S3), reflecting the low dimensionality of the underlying code (here, Dim *<* 4 for each rank, computed using the methods of (Recanatesi et al., 2022)).

In summary, this simulation demonstrates that inferential planning dynamics reproduce the neural population (information) geometry observed by (Xie et al., 2022) during the delay period, when monkeys maintain a sequential plan for three saccades in working memory.

### 2.4 Inferring and holding in memory a variable-length sequence of forward or backward saccades

Third, we consider an experiment (Chen et al., 2024) with an identical setup to the previous task but extended to three novel conditions (Figure 4A-B and Figure 5A-B). In the first (Forward) condition, monkeys observed sequences of *variable length*—from one to three targets—and therefore had to infer the length of their plan. In the second (Backward) condition, monkeys again observed sequences of variable length but were required to reproduce the plan in the backward direction, that is, to make saccades to the targets in the reverse order of presentation. Finally, in the third (Mixed) condition, monkeys observed sequences of fixed length (two targets) and then received a cue instructing them to reproduce the plan either in the forward direction (the same order of presentation) or in the backward direction (the reverse order).

**Figure 4.**
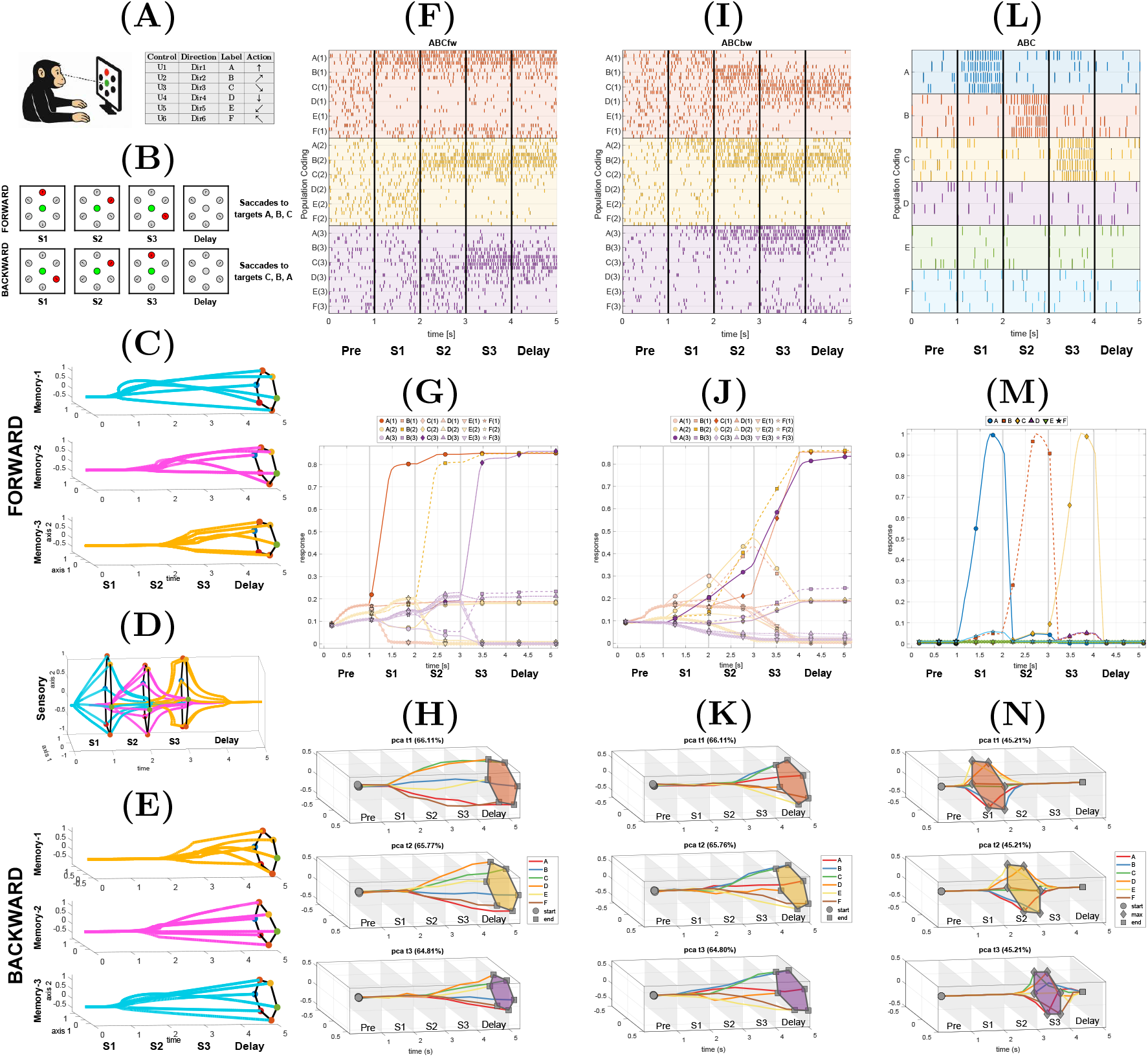
Inferring and holding in memory a variable-length sequence of saccades, to be executed in either the forward or the backward direction. This simulation reproduces the first two conditions of the experimental setup of (Chen et al., 2024). In (A-B), sequences of variable length (one to three elements) are drawn from six possible target {*A* –*F*} (e.g., *ABC, AB, ABD, BCA, CB, A, B*). For each block of trials, the instruction is to later execute the trial in the *forward* or *backward* direction. The targets are presented at stages S1, S2, and S3 (but S2 and S3 targets can be omitted). During the subsequent delay period, the inferred sequence is maintained in working memory. (C-E) Schematic illustration of the results reported in (Chen et al., 2024) showing neural state activities corresponding to forward ‘memory’ populations, ‘sensory’ populations, and ‘backward’ memory populations. (F-H) Simulation of the *ABC* sequence in the Forward condition. (F) Raster plots showing simulated neural activity, with four neurons coding for each of the 6 control states (Targets *A* −*F*) at Level 2, at each each of the three time steps. Neurons coding for control states at the 3 time steps are colour coded in red, yellow and violet, respectively. At the end of the simulation, corresponding to the delay period before execution, the policy *ABC* is coded by the simultaneous activation of three populations of neurons, *A*(1), *B*(2), and *C*(3). (G) The same results, but plotted as simulated firing rates of neurons associated with different actions. (H) The same results, but plotted as simulated dynamics of the first three principal components (PCs) summarizing the neural state space shown in (C) and corresponding to the ‘memory’ subspaces in the forward case in (Chen et al., 2024). (I-K) Simulation of the *ABC* sequence in the Backward condition, using the same format as (F-H). These activations summarize neural state space shown in (E) and corresponding to the ‘memory’ subspaces in the backward case in (Chen et al., 2024). (L-N) Simulation of Level 1 activations during the observation of the *ABC* sequence in both the Forward and Backward conditions. This simulation follows the same format as (F–H) but depicts the activity of hidden states at Level 1 rather than Level 2. These activations summarize the neural state space shown in (D) and corresponding to the ‘sensory’ subspaces in (Chen et al., 2024). See the main text for further explanation.

**Figure 5.**
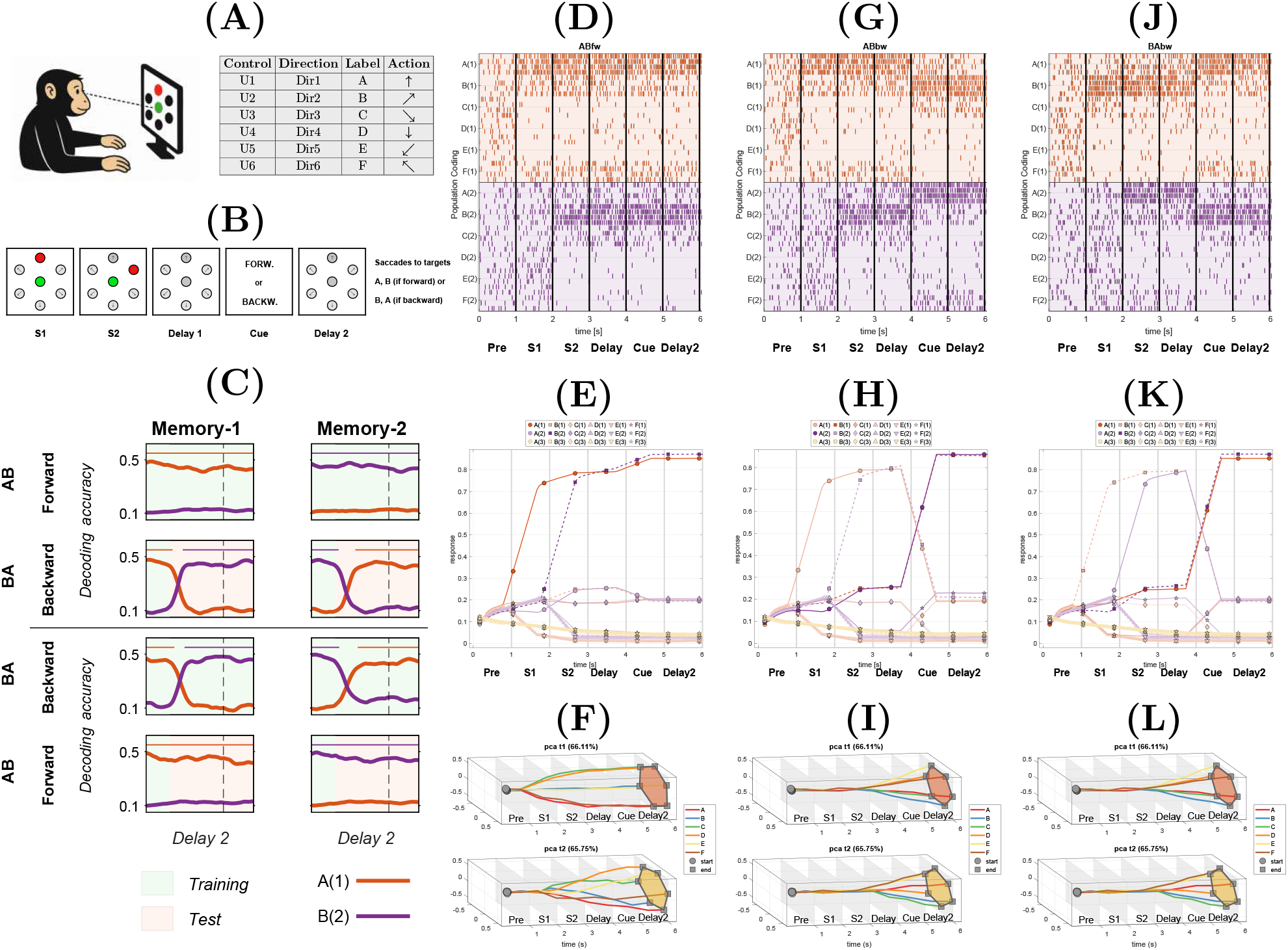
Inferring and holding in memory a fixed-length sequence of saccades, to be executed in either the forward or backward direction. This simulation reproduces the third condition of the experimental setup of (Chen et al., 2024). (A-B) In this task, the sequence length is fixed to two targets, selected from the set of six possible targets {*A*− *F*} . The two targets are presented sequentially during epochs *S*1 and *S*2, followed by a delay period and an instruction cue indicating whether the sequence should be reproduced in the *forward* or *backward* direction. During the subsequent delay period, the inferred plan is maintained in working memory. (C) Figure adapted from (Chen et al., 2024), showing that the first ‘mem-ory’ subspace (‘memory-1’) in the forward condition could be generalized to the second ‘memory’ subspace (‘memory-2’) in the backward condition, and the second ‘memory’ subspace (‘memory-2’) in the backward condition could be generalized to the first ‘memory’ subspace (‘memory-1’) in the forward condition. The horizontal coloured bars indicate time windows where the accuracy of an algorithm trained to generalize from forward to backward ‘memory’ subspaces (or vice versa) was significantly higher than the chance level, indicating that the subspaces could be effectively generalized. Training and test epochs for the algorithm are shown in green and orange, respectively. (D-F) An example problem: inferring the *AB* sequence in the forward condition. (D-E) Raster plots showing simulated neural activity and simulated firing rates associated with inferring the *AB* plan. (F) Simulated dynamics of the two first principal components during the task. (G-I) Another example problem: inferring the *AB* sequence in the backward condition. (J-L) Another example problem: *BA* sequence in the backward condition. See the main text for further details.

As in (Xie et al., 2022), the first, second, and third actions in the sequence were represented in separate subspaces—here referred to as ‘memory’ subspaces. Additionally, the dynamics of these subspaces exhibited a gradual build-up of activity at the moment when the animal could infer its position within the sequence. For example, if the animal knew that the sequence had to be executed in the forward direction and observed target A at ‘S1’ it could already infer A(1), even if the actual sequence length was still unknown. This was reflected in a rapid build-up of activity corresponding to A at ‘S1’ (Figure 4C). Conversely, if the animal knew that the sequence was to be executed in the backward direction and observed target A, it could not yet infer whether the correct action was A(1), A(2), or A(3), resulting in no such build-up of activity (Figure 4E). Furthermore, during observation of the three targets (S1, S2, S3), additional ‘sensory’ subspaces transiently coded for the observed targets (Figure 4D). Finally, an analysis of the third (Mixed) condition revealed that monkeys used common subspaces across both forward and backward tasks: the first ‘memory’ subspace (‘memory-1’) in the forward condition could be generalized to the second ‘memory’ subspace (‘memory-2’) in the backward condition, and the second ‘memory’ subspace (‘memory-2’) in the backward condition could be generalized to the first ‘memory’ subspace (‘memory-1’) in the forward condition (Figure 5C).

We used the same hierarchical generative model as in the previous simulation, extended to include task-rule information (‘forward’ vs. ‘backward’) at Level 2. This rule information is represented in two ways. First, Level 2 contains a hidden state (‘context’) that encodes which rule is currently in effect. In the first two conditions, this context is known in advance of observing the target sequence, whereas in the third condition it must be inferred when the forward/backward cue is presented. Second, Level 2 includes two rule-specific sets of policies, one for the forward rule and one for the backward rule. These different policies can generate the same observable action sequences. For example, the sequence *AB* can arise either from a forward policy *AB* or from a backward policy *BA*. Although these policies differ in their structure, they share the same action components, *A*(1) and *B*(2). Policy selection between forward and backward alternatives is driven by the inferred context, which combines prior beliefs with available task cues.

Figure 4F-N shows the simulation of the first two (Forward and Backward) conditions of the (Chen et al., 2024) study, in which the monkeys know from the start whether the plan will be executed in the forward or backward direction but do not know the sequence length. Figure 4F-H shows the simulation of the Forward condition. The gradual build-up of neural activity associated with the *ABC* sequence to be executed in the forward direction can be seen at three different levels: in the simulated raster plots (Figure 4F), the simulated firing rates (Figure 4G), and the simulated ‘memory slots’, which simply correspond to 3D projections of the principal components (PCs) of simulated neural activity during inference (Figure 4H). In all cases, the build-up of the first target in the plan (A(1)) is faster than that of the second (B(2)) and third (C(3)) targets, consistent with the experimental data (Figure 4C). See also Supplementary Materials, Figure S3, for an illustration of the case of a sequence of length two.

Figure 4I-K presents the results of the simulation for the Backward condition. The sequence of target presentations (*A, B*, and *C*) was identical to that in the Forward condition, but the temporal dynamics of activity differed. Following the presentation of the first two targets (*A* and *B*), the model maintained multiple concurrent hypotheses about their possible positions in the sequence (*A*(1), *A*(2), *A*(3), *B*(1), *B*(2), *B*(3)), reflecting uncertainty about sequence length. This ambiguity persisted until the third target (*C*) was observed, at which point its identity as the final element (*C*(3)) became unambiguous, enabling rapid resolution of uncertainty and selective build-up of activity corresponding to the third step. This pattern parallels the empirical findings (Figure 4E).

Additionally, Figure 4L-N illustrates the transient activation of Level 1 actions inferred during the observation of the three targets, corresponding to the ‘sensory subspaces’ illustrated in Figure 4D. Within the hierarchical model, these Level 1 activations arise from bottom–up inference of the currently observed targets and provide evidence that updates higher-level (Level 2) beliefs about the sequence. Unlike the sustained Level 2 representations, which encode abstract ‘memory’ subspaces, Level 1 activations are short-lived and confined to the sensory epochs, reflecting the transient message passing required for hierarchical inference.

Figure 5D-L shows the simulation of the third condition of the study of (Chen et al., 2024), in which monkeys know from the beginning that the plan consists of two targets but are informed about the forward or backward direction only when the cue is presented. In this case, after observing two targets, *A* and *B*, the monkeys’ neural population dynamics reflect a higher probability of the “forward” (*AB*) plan (Figure 5C, see also Tian et al., 2024). In our model, this happens when the animals assign a higher a priori probability to “forward” compared to “backward” policies, reflecting their “natural statistics” of experience – and the fact that series of events typically unfold in the forward direction (Without such prior, after observing the two targets, *A* and *B*, the model would maintain parallel, equally plausible, hypotheses, *A*(1), *A*(2), *B*(1), and *B*(2), see Supplementary S4). After receiving a ‘forward’ cue, the model increases its confidence that the correct plan is *AB* (Figure 5D-F). Conversely, after receiving a ‘backward’ cue, the model reverses its initial hypothesis and correctly infers the *BA* plan (Figure 5G-I). Similarly, after observing the two targets, *B* and *A*, and a ‘backward’ cue, the model correctly infers the *AB* plan (Figure 5J-L). Crucially, the model reuses common ‘memory subspaces’ for the first and second targets across both forward and backward tasks, as observed empirically (Figure 5C). For example, the neural population encoding *A*(1) is the same whether the model has observed targets *A* and *B* followed by a ‘forward’ cue (Figure 5A), or targets *B* and *A* followed by a ‘backward’ cue (Figure 5G).

In summary, these simulations illustrate how inferential planning dynamics reproduce the near-orthogonal, hexagon-shaped subspaces for sequential targets, which are shared across forward and backward tasks, as reported by (Chen et al., 2024).

## 3 Discussion

In this paper, we have shown that the recently discovered ‘activation slots’ are an effective neural substrate for inferring arbitrary plans. Further, we showed the activity of model neurons during inference match experimentally recorded neurons in a variety of planning and sequence memory tasks (Mushiake et al., 1991; Xie et al., 2022; Chen et al., 2024). We anticipate our planning as active inference formulation, using message passing between activation slots, will serve as a framework for future fine grained mechanistic understanding of planning in prefrontal cortex (and beyond) at the level of neuron and synapse.

Indeed, concurrent modelling work has shown that when RNNs are trained on spatial planning tasks, they learn to plan with activation slots—with plans as fixed points of attractor dynamics—with the synaptic connections between slots mirroring the connectivity structure of the spatial environment (Jensen et al., 2025). This bears a relation to our inferential planning via message passing between slots (where, technically, the attractors are variational free energy minima). Our inferential planning approach, however, not only provides a formalism for *arbitrary* planning tasks, but also provides a formal answer to a key question about neural representation: why plan elements are coded simultaneously. In inferential planning (e.g., active inference, planning as inference, and related approaches (Attias, 2003; Botvinick & Toussaint, 2012; Lázaro-Gredilla et al., 2024; Friston et al., 2017a; Parr et al., 2022; Ortega & Braun, 2013; Gershman & Beck, 2017; Kappen et al., 2012; George et al., 2021; Pezzulo et al., 2018; Levine, 2018; Isomura et al., 2022; Bastos et al., 2012)), a plan is generated by inferring the most likely sequence of actions (or policy) leading from initial to goal states. Crucially, this inferential process entails reciprocal message passing among representations of past, present, and future (expected) states, which must therefore be maintained and updated in parallel (George & Hawkins, 2009; George et al., 2025; Friston et al., 2017a,b). The simultaneous coding of plan elements is therefore not a nuance but a necessary prerequisite for the message passing that underlies planning.

Other models (beyond the aforementioned RNNs (Whittington et al., 2025; Jensen et al., 2025)) also represent all elements in a sequence simultaneously. Competitive queuing models (Bullock, 2004) generate serial order via a competitive process in which the most active unit wins the competition and generates the corresponding action; it then inhibits itself, allowing the next most active unit to drive the subsequent action, and so forth. This differs from our framework which can endogenously generate a sequence of plan element activations without requiring a predefined queuing mechanism. Indeed, in Figure 2F,I, the firing rate of the first cursor movement increases faster than that of the second and third ones, reflecting a faster resolution of uncertainty about what to do next. The neural network model of (Botvinick & Watanabe, 2007) introduced the idea that prefrontal neurons form conjunctive representations of item identity and temporal order through gain modulation, analogous to our Level 2 neural populations, which selectively encode both state (or action) identity and temporal position. However, that model was primarily developed to explain the representation of serial order in working memory, whereas our framework addresses how sequential plans are inferred and maintained. Other models represent multiple ordered elements of a sequence simultaneously along a ‘mental line,’ enabling transitive inference (Jensen et al., 2015; Di Antonio et al., 2024; Mannella & Pezzulo, 2025). This contrasts with our framework, which uses simultaneous representations of actions and states to construct and update plans, rather than to encode relational order.

While building an understanding—using a probabilistic formalism—is one level abstracted from neuron and synapse, it will not only be helpful when interpreting representations from tasks that manipulate transition probabilities between goals or those with cue uncertainty (Findling et al., 2025), but also provides a formal account of any neural representation that corresponds to a variable in the underlying generative model. Some of our design choices—such as cosine tuning (or linear mixed selectivity)—are made to match empirical data, demonstrating that this modelling approach can accommodate such findings, though not that they are necessary consequences of it. Other aspects, however, such as the duplication of actions across subspaces (i.e., nonlinear mixed selectivity), are not optional but required for inferential planning. Our model therefore offers a novel perspective on the empirically reported orthogonal coding of subspaces, suggesting that the duplication of actions across subspaces may support the message passing required for inferential planning, rather than serving solely a working-memory function. Furthermore, the inferential dynamics illustrated in our simulations emerge from message passing among neural populations and are not “baked into” the model. Notably, specific features of these dynamics—such as the order in which actions are inferred, exemplified by the sequential inference of the first action before the second and third in the simulation of (Mushiake et al., 2006)—arise naturally from the inferential process and can be directly related to empirical observations. Furthermore, our probabilistic framing suggests that the ‘sensory subspaces’ observed in (Chen et al., 2024) correspond to Level 1 representations that play a central role in hierarchical inference, rather than serving merely as temporary storage. This could be tested by perturbing these sensory subspaces (e.g., optogenetically), which should disrupt the animal’s ability to correctly infer targets, as shown in other studies using hierarchical inference (Proietti et al., 2023; Van de Maele et al., 2024; Donnarumma et al., 2025). Our framework also suggests that, in sequential tasks such as those studied by (Xie et al., 2022; Chen et al., 2024), neural dynamics may encode implicit expectations about future targets even before they are observed. One example can be appreciated in the raster plot of Figure 3F. In the third column (period *S*2), after observing *A*(1) and *B*(2), the activity associated with *A*(3) and *B*(3) decreases, reflecting the fact that targets in this paradigm are sampled without replacement. In contrast, the activity associated with *C*(3), *D*(3), *E*(3), and *F* (3) increases, reflecting the expectation that one of these targets will appear next. Furthermore, the framework suggests that the ‘reversals’ from forward to backward plans observed following a backward cue in (Chen et al., 2024; Tian et al., 2024) may emerge through probabilistic belief updates, whereby initially favored forward policies are replaced by backward policies inferred from the cue. The empirical validity of these signatures of inferential dynamics remains to be assessed in future experimental work.

There are several (addressable) limitations of our model. First, we illustrated our results using a specific factorization of the generative model, combining linear and nonlinear mixed selectivity, which captures key empirical observations such as the presence of separable, low-dimensional, and near-orthogonal subspaces in frontal cortex. However, we deliberately kept this factorization relatively simple compared to experimental observations in (Xie et al., 2022; Panichello et al., 2024), which suggest that these subspaces may be distributed across larger and more overlapping neural populations. This choice reflects the primary aim of our study, which was not to propose a detailed account of the neural coding of subspaces, but rather to investigate how separable neural representations can support inferential planning of sequential actions. The inferential framework itself is compatible with alternative factorizations, including different degrees of mixed selectivity or continuous tuning profiles (e.g., cosine tuning). Future work could attempt to infer the factorization and dimensionality of the hidden states of the generative model directly from neural recordings in a more bottom-up manner.

Second, we provided simplified simulations of the neural dynamics associated with inference. While these simulations are sufficient to support our main points—such as increases (or decreases) in population activity associated with inferring a particular action at a given time point—they omit additional details, including the precise shape of ramping activity that may reflect increasing confidence in the inferred plan over time, as well as biological details of single-cell dynamics (e.g., refractory periods).

Third, we only analysed the planning phase, rather than the execution phase. Thus our neural subspaces correspond to distinct (and fixed) positions (e.g., 1,2,3,…) of the plan. This is subtly different to the slots model (Whittington et al., 2025) in which slots correspond to relative positions (e.g., present, past, or future) with slot contents moving between slots so that the present slot always contains the present sequence element. This relative representation makes readout (i.e., execution) extremely efficient as there only needs to be one set of readout weights. While we modelled planning with fixed slots, we posit that, in prefrontal cortex, during the execution phase the contents of slots will shuffle between one-another to take advantage of the simple readout mechanism. Indeed experimental data suggests this, with planning phases using using fixed slots (Chen et al., 2024) (like our model), but execution passing contents between slots (El-Gaby et al., 2024) (like the slots model).

Fourth, for simplicity, our simulations modeled only a discrete set of cursor movements and saccade directions. To reproduce behavioral dynamics more faithfully, the discrete framework presented here could be extended by coupling it with a continuous-time model of cursor or eye movements, yielding a hybrid discrete–continuous active inference architecture (Parr & Friston, 2018b; Priorelli et al., 2025).

Frontal cortex has long been known to be central to higher level cognitive functions including cognitive control (Miller & Cohen, 2001; Duncan, 2001; Pezzulo et al., 2018; Koechlin et al., 2003; Barceló, 2021; Proietti et al., 2025), cognitive map formation (Schuck et al., 2016; Wang & Hayden, 2021; Behrens et al., 2018), working memory (Curtis & D’Esposito, 2003) and planning (Mattar & Lengyel, 2022; Koechlin, 2016; Goel & Grafman, 1995; Duncan et al., 1996). In this work we provide a formalism of planning built from recent experimental results for simultaneous sequence element coding in prefrontal cortex. In doing so we provide explanations for several empirical observations, mechanistically link planning and working memory, and generating testable predictions for future experiments.

## 4 Methods

In this section, we first provide a concise formal overview of the active inference framework that we adopt in this study (Parr et al., 2022) (Section 4.1). Next, we describe the message-passing scheme that underlies neural simulations in active inference (Section 4.2) and detail the specific hierarchical generative model used to implement the simulations (Section 4.3). Finally, we outline the analytical procedures employed to characterize low-dimensional neural subspaces in the simulation of the study by (Xie et al., 2022; Chen et al., 2024) (Section 4.5).

### 4.1 Brief introduction to Active Inference

Active Inference provides a normative account of perception–action as *variational Bayesian inference* under a generative model, coupled with *policy selection* that minimizes expected free energy (EFE) (Parr et al., 2022). An agent maintains a probabilistic model over latent (hidden) states and observations—typically represented as a probabilistic graphical model (Bishop, 2006)—and acts to realize outcomes that jointly pursue utility maximization (*pragmatic value*) and uncertainty minimization (*epistemic value* or *information gain*). In the following, we summarize the key ingredients of active inference, from model formalization to the steps required to perform inference by updating hidden states, policies, and precision.

#### Generative model

Let *O*_0:*T*_ = (*O*_0_, …, *O*_*T*_) denote a sequence of *T* observations, *S*_0:*T*_ a sequence of *T* hidden states, *π* = (*π*_1_, …, *π*_*T*_) a policy (a sequence of control states, or more simply “actions”), *γ* ∈ ℝ_+_ a precision controlling the sharpness (i.e., inverse temperature) of policy selection, and **Θ** the model parameters. A convenient factorization (here, dealing with a single hierarchical level) is:

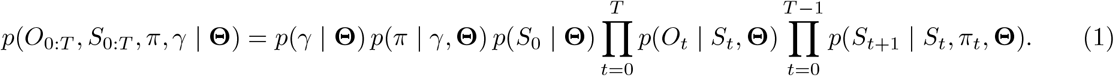

#### Parameterization

We write **Θ** = {**A, B, C, D, E**, *β*} with:

- **Likelihood A**: *p*(*O*_*t*_ | *S*_*t*_, **Θ**) = **A**. It maps hidden causes *S* to observations *O* (often categorical).
- **Transitions B**: *p*(*S*_*t*+1_ | *S*_*t*_, *π*_*t*_, **Θ**) = **B**(*π*_*t*_), i.e., policy-conditioned dynamics.
- **Preferences C**: an *a priori* (log-)distribution over outcomes, encoding what the agent prefers to observe, *P* (*O*_*τ*_) ≡ **C**.
- **Initial prior D**: prior on hidden states *p*(*S*_0_ | **Θ**) = **D**.
- **Policy prior E**: habitual/structural prior over policies.
- **Precision** *γ*: sampled from a Gamma prior *p*(*γ* | **Θ**) ∼ Γ(1, *β*).

Intuitively, **A** tells the agent ‘what sensations to expect from each state’; **B** tells it ‘how hidden states change under a chosen policy’; **C** encodes preferences; **D** is what it believes before seeing anything; **E** captures habits. To move to a hierarchical specification, we would condition the **D** hidden state priors for one level on the hidden states at the level above, such that a hidden state at any given level of the model predicts the initial state of a short sequence at the level below.

#### Approximate posterior and mean-field form

A common implementation of active inference uses a tractable variational posterior (see mean-field theory Parisi, 1988):

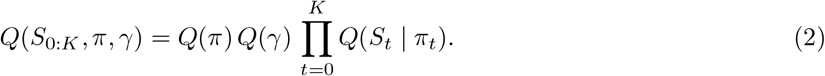

With this distribution, we maintain a posterior over the policy *Q*(*π*) and, for each time step, a posterior over hidden states conditional on the policy.

#### Variational free energy (VFE)

Perceptual inference at time *t* minimizes

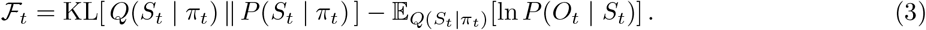

The first term is the Kullback-Leibler (KL) divergence representing the *complexity* cost (deviation from prior), the second is the expected negative log-likelihood of the observation (negative *accuracy*, lack of fit to data). Balancing the two yields predictive beliefs (Parr et al., 2022).

Under a standard categorical parameterization, a fixed-point update has the form

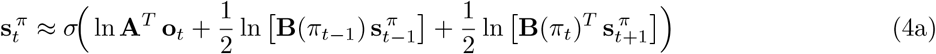

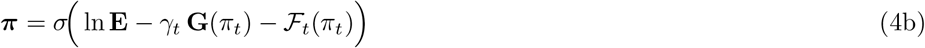

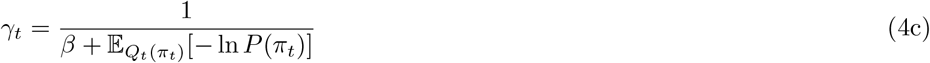

Here *σ*(·) is the componentwise Softmax, 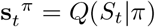 denotes the posterior state marginals and **o**_*t*_ is the one-hot encoding of the observed outcome at time *t, Q*(*π*) = Cat(***π***) is the posterior over *π*, while *Q*_*t*_(*π*_*t*_) and 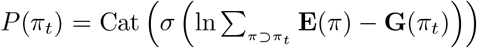 are respectively the posterior and the prior over *π*_*t*_. Intuitively, Eq. (4a): this combines current sensory evidence with predicted messages that depend upon predictions based on beliefs about previous states and postdictions. Note that in Eq. (4a), **B** and **B**^*T*^ are both assumed to have normalized (sum-to-one) columns. from beliefs about subsequent states; Eq. (4b): prefer policies that are *habituated* (large ln **E**) and have *low* EFE; Eq. (4c): increase precision when expected free energy is low/consistent. The form of Eq. (4a) depends upon a marginal approximation to the prior for states in Eq. (3) that accounts for beliefs about both past and future states (c.f., Bayesian smoothing). For technical details of this scheme, please see (Parr et al., 2019a) and the Appendices of (Parr et al., 2021; Friston et al., 2017a).

#### Expected free energy (EFE) and action selection

Policies are scored by their EFE:

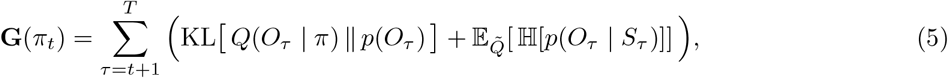

where 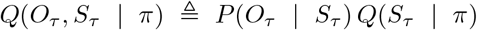 is the predictive posterior distribution, 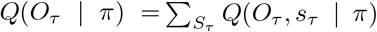 is the predicted outcome, *P* (*O*_*τ*_) ≡ **C** encodes preferences, and ℍ [·] is Shannon entropy. Intuitively, the first term (*expected cost* /risk) favours policies that make preferred outcomes likely; the second (*expected ambiguity*) favours policies that reduce uncertainty by seeking informative states.

In this model, action selection corresponds to the generation of predicted control signals, which are sampled from the policy posterior in Eq. (4b). These samples represent planned actions, reflecting the agent’s internal simulation of future behaviour under each candidate policy (Parr et al., 2022).

### 4.2 Message Passing and neural architecture

In this section, we briefly outline the way in which we can interpret the fixed point scheme outlined about in terms of neuronal message passing, with a focus on Eq. (4a). The idea is relatively simple. If we assume neuronal membrane potentials for a pool of neurons, on average, are represented by a vector 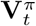, and firing rates (again, on average) by 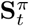, then treating the softmax function as a neuronal transfer function, such that 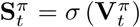, we arrive at an interpretation (up to an additive constant) of 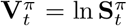. We can then express the dynamics of the membrane potentials to a first order approximation as a simple attractor system whose fixed point corresponds to the solution of Eq. (4a):

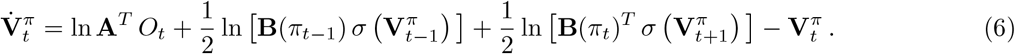

Here, the key points to note are that the membrane potentials associated with a given population (indexed by time and policy) that represent beliefs *about* a particular time evolve such that they depend upon the potentials of populations representing beliefs about the immediate past and future. This implies that at any given time, we need to simultaneously hold beliefs *about* other times to instantiate these dynamics. It is this distinction between the time at which beliefs are held, and the times those beliefs are *about* that underwrites the core ideas in this paper.

Before we move on, it is worth highlighting that this property is not specific to the message passing scheme outlined here, and would also apply to strict belief-propagation or variational message passing schemes. The reason for this is inherent in the structure of the generative model, rather than of the particular method of solution. Specifically, the model used here relies in part upon a Markov chain, in which the Markov blanket of any given state includes both its predecessor and successor. As such, beliefs about proximal time-points will always be informative about the current time.

### 4.3 Hierarchical generative model for planning

The hierarchical Active Inference model employed in this study is designed to structure motor planning across two hierarchical levels, denoted by *l*. At each level, the model encodes hidden variables that represent distinct aspects of motor organization: low-level motor actions at Level 1 and structured sequences of these actions at Level 2. The overall architecture of the model is illustrated in Figure 6.

**Figure 6.**
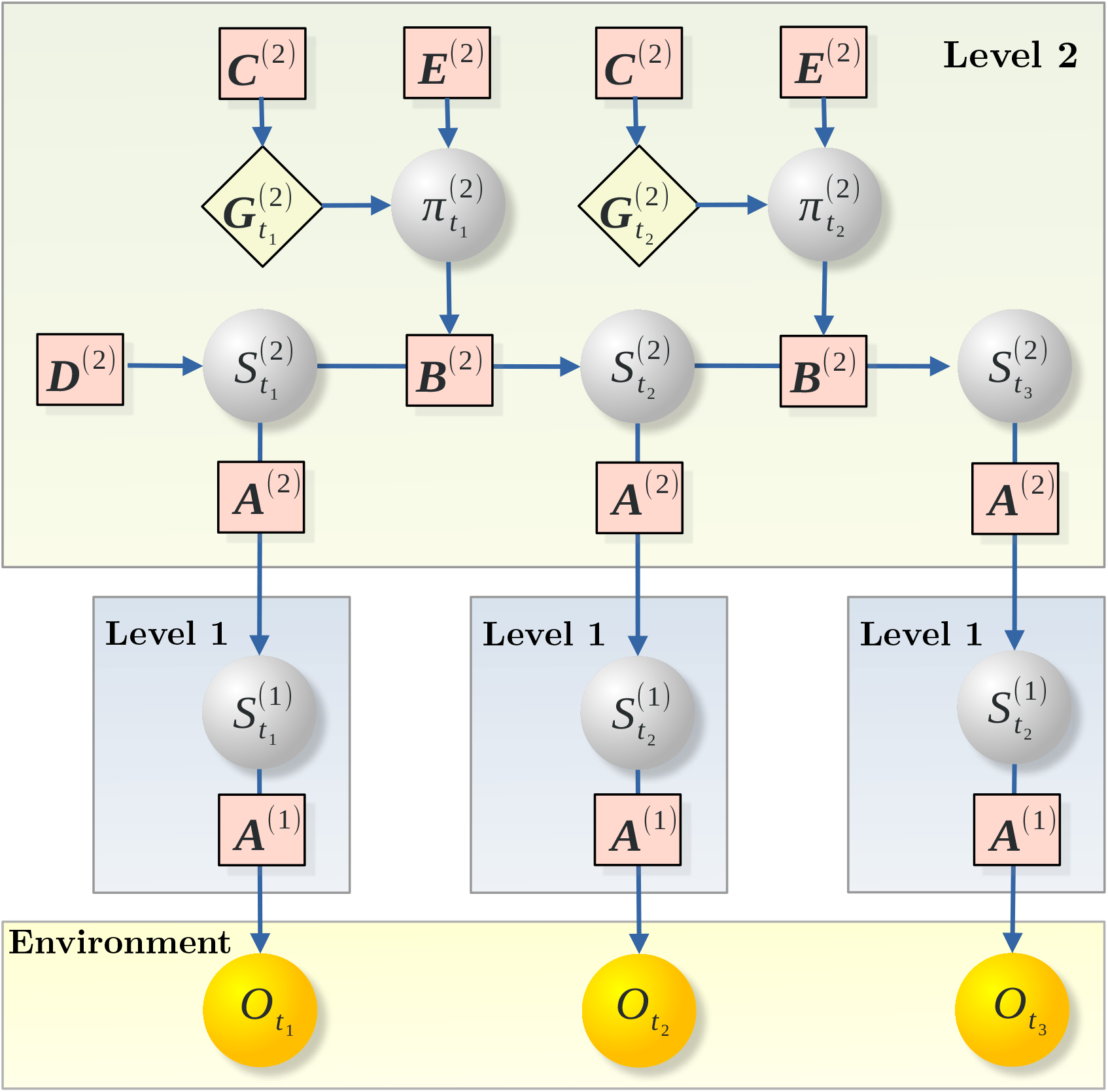
The hierarchical (deep) Active Inference model for action planning. The model consists of two interacting layers, Level 1 and Level 2, organized within the Active Inference framework. Gray nodes denote hidden states and policies, yellow nodes represent observations, and edges illustrate probabilistic dependencies between variables. Note that although the edges are unidirectional, reflecting probabilistic dependencies, in the generative model information flow is effectively bidirectional, reflecting reciprocal message passing between neural populations across levels. At the higher level, policies *π* encode sequences of control states (abstract actions), whereas at the lower level, they correspond to single motor commands. Rectangular nodes depict probabilistic distributions – the conditional dependencies defining the Hidden Markov Models - parameterized by the matrices **A, B, C, D**, and **E**. The **A** matrix encodes the likelihood model (how hidden states *S*_*t*_ generate observations *O*_*t*_); the **B** matrix parameterizes state transitions under each policy *π*; the **C** matrix specifies preferred outcomes and contributes to the expected free energy *G*; the **E** matrix represents prior preferences over policies (representing habits). Notably, the hidden states 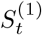 at Level 1 serve as *observations* for the higher-level process at Level 2, linking temporal inference across hierarchical timescales. See the main text for further explanation.

To capture the structured dependencies among motor components, the hidden state space at each level *l* is expressed as a tensor product:

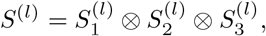

where each factor (subscripts here being factor indices, and not time-points as previously) encodes a specific dimension of motor planning and control:

- **Hidden Control States** 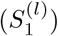 – These variables specify the motor controls of the actions themselves. At the lowest level (*l* = 1), they represent discrete atomic actions, while at the higher level (*l* = 2) they encode sequences of actions composing a coherent motor strategy.
- **Hidden Location States** 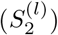 – This factor encodes the spatial component of the motor plan. At Level 1, it corresponds to concrete movement execution in physical space, whereas at Level 2, it represents the structural organization of sequence of action locations corresponding to the policy.
- **Hidden Context States** 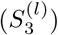 – This component defines the contextual or temporal structure of the motor sequence. In our simulations, at Level 2 it specifies the *order* in which actions are to be planned (e.g., forward vs. backward order relative to the presentation).

#### Observations

Each observation *O* = *O*_1_ ⊗ *O*_2_ ⊗ *O*_3_ is defined as the tensor product of the following components:

- *O*_1_: the observation of control actions.
- *O*_2_: the observation of target locations.
- *O*_3_: the cue indicating the execution order, which specifies the sequence in which actions should be performed.

The cue *O*_3_ is used in Simulation (4) (see Table 1) and includes options such as ‘forward,’ ‘backward,’ other possible execution orders, and ‘idk’—a neutral cue indicating that execution does not depend on the presentation order.

**Table 1.** Schematics of the simulation parameters. The table summarizes hierarchical Active Inference settings for four tasks: (1) planning sequences of cursor movements ((Mushiake et al., 2006), Fig. 2); (2) inferring and holding in memory a sequence of saccades ((Xie et al., 2022), Fig. 3); (3) inferring and holding in memory a variable-length sequence of forward or backward saccades ((Chen et al., 2024), Fig. 4); and (4) inferring and holding in memory a fixed-length sequence of saccades executed in either forward or backward order ((Chen et al., 2024), Fig. 5). For each task, the table reports hidden state configurations, prior preferences, and policy constraints. Simulation (1) uses environment-constrained policies and goal preferences. Simulation (2) expands target states across time steps to encode temporal information, with no contextual cue. Simulation (3) introduces variable-length sequences with contextual cues known from the start (Forward or Backward). Simulation (4) uses fixed-length sequences with preferences for both directions, where the cue is revealed at time step *t*_cue_.

| Task | Settings | Description |
| --- | --- | --- |
| Simulation (1):<br>Planning sequences of<br>cursor movements<br>(Mushiake et al., 2006),<br>Fig. 2) | $S_1^{(2)} \in \{\text{Up, Down, Left, Right, Stay}\}$<br>$S_2^{(2)} \in \{S_{11}, S_{12}, \dots, S_{55}\}$<br>$C_2^{(2)} = \text{Goal}$<br>$C_3^{(2)} = \emptyset$<br>$\mathbf{E}^{(2)} = p(\pi \mid \text{Maze})$<br>$T^{(2)} = 3$ | 5 cursor controls<br>25 target locations<br>Preference on goal state<br>No contextual cue<br>Prior constrained by environment<br>Fixed-length sequences (3) |
| Simulation (2):<br>Inferring and holding<br>in memory a sequence of<br>saccades (Xie et al., 2022),<br>Fig. 3) | $S_1^{(2)} \in \{A, B, C, D, E, F, \text{Stay}\}$<br>$S_2^{(2)} \in \bigcup_{t=1}^T \{A(t), \dots, F(t), \text{Empty}(t)\}$<br>Sequence Length = 3<br>$\mathbf{C}_3^{(2)} \in \{\text{Forward}\}$ | Six directions + no move<br>targets + empty for each step<br>Fixed-length sequences (3)<br>Contextual cue known from start |
| Simulation (3):<br>Variable-length sequence<br>of forward/backward saccades<br>(Chen et al., 2024),<br>Fig. 4) | $S_1^{(2)} \in \{A, B, C, D, E, F, \text{Stay}\}$<br>$S_2^{(2)} \in \bigcup_{t=1}^T \{A(t), \dots, F(t), \text{Empty}(t)\}$<br>$\mathbf{C}_3^{(2)} \in \{\text{Forward, Backward}\}$<br>Sequence Length $\in \{1, 2, 3\}$ | Six directions + no move<br>targets + empty for each step<br>Contextual cue known from start<br>Variable-length sequences |
| Simulation (4):<br>Fixed-length sequence with<br>forward/backward execution<br>(Chen et al., 2024),<br>Fig. 5) | $S_1^{(2)} \in \{A, B, C, D, E, F, \text{Stay}\}$<br>$S_2^{(2)} \in \bigcup_{t=1}^T \{A(t), \dots, F(t), \text{Empty}(t)\}$<br>Sequence Length = 2<br>$O_3(t_{\text{cue}}) \in \{\text{Forward, Backward}\}$ | Six directions + no move<br>targets + empty for each step<br>Fixed-length policies (2)<br>Contextual cue observed |

#### Transition mapping

The transition probability 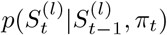 given control states is specified by the tensor:

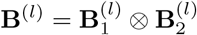

where:

- 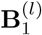 governs transitions among control states.
- 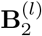 defines transitions among spatial locations across time steps.

Transitions are nearly deterministic, but stochasticity is introduced to model execution errors or uncertainty in control signals.

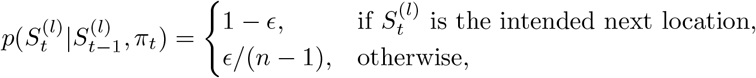

where *ϵ* is a small noise term and *n* is the number of allowable possible locations (where a transition is actually possible).

#### Low-level generative model (Level 1)

In simulations of tasks from (Mushiake et al., 2006), 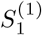 represents the hidden controls of the cursor actions (i.e. Up, Down, Left, Right). In simulations of tasks from (Xie et al., 2022) and (Chen et al., 2024), 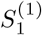 represents the hidden control for the saccadic movements towards the targets A, B, C, D, E, F. In both cases, 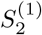 represents the hidden locations of the target states.

#### Likelihood mapping at Level 1

The likelihood *p*(*O*^(1)^ | *S*^(1)^) between hidden states *S*^(1)^ and observations *O*^(1)^ is specified through the tensor:

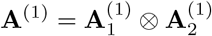

where:

- 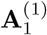 maps hidden control states 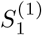 to observations *O*_1_ (primitive control actions).
- 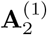 maps hidden target states 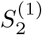 to observations *O*_2_ (target locations).

The mapping is almost deterministic but implements *cosine tuning* for sensory encoding:

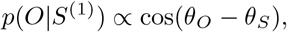

where *θ*_*O*_ and *θ*_*S*_ represent preferred directions of observation and hidden state, respectively. Recognition accuracy is tuned in order to get approximately 85% with respect of the preferred direction, and misclassification occurs toward the closest state in angular space.

#### Higher-level generative model (Level 2)

<TC-F-BlockquoteIndent>Level 2 comprises three hidden-state factors, analogous to those in Level 1, representing beliefs about *Controls* 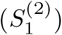 and *Locations* 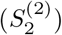. The third factor, *Context* 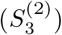, encodes the sequence execution order (e.g., ‘forward’, ‘backward’) or its absence (‘idk’). This contextual factor may be fixed prior to the trial (as in simulations from (Xie et al., 2022)) or revealed during the trial through observation of the cue *O*_3_ (as in simulations from (Chen et al., 2024)).

#### Likelihood mapping at Level 2

The likelihood *p*(*S*^(1)^ |*S*^(2)^) specifies how higher-level hidden states *S*^(2)^ predict lower-level hidden states *S*^(1)^. It is represented by the tensor:

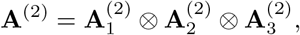

where:

- 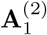: maps higher-level (sequence of) control states 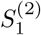 to lower-level controls 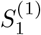;
- 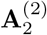: maps higher-level (sequence of) target states 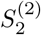 to lower-level locations 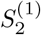;
- 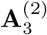: maps higher-level context states 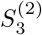 to the execution order cue *O*_3_.

Here, higher-level states are mapped onto lower-level states (or cues in the case of *S*^(^2)_3_, which are treated as observations in the hierarchical model. Formally, the likelihood mappings are defined as 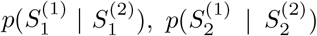, and 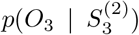, with each mapping being nearly deterministic while allowing for a small degree of stochasticity (approximately 5% noise) in recognizing the correct correspondence. We imposed cosine directional tuning (a form of linear mixed selectivity) on Level 2 states by specifying, in the likelihood matrix, a strong probabilistic mapping (0.85) between each hidden state and its preferred direction (e.g., “target A”), together with progressively weaker mappings to neighboring targets as angular distance increased (weaker for “B” and “F”, weaker still for “C” and “E”, and weakest for “D”); see Section 4.6 for details.

Observations *O*_3_ (e.g., ‘Forward’ or ‘Backward’ cues) are included in tasks from (Chen et al., 2024) to indicate execution order.

#### Preferences

The tensor 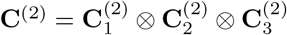 encodes prior preferences over observations:

- 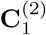: preferences over motor actions 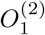 (goal-directed behaviour).
- 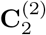: preferences over spatial positions 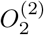 (preferred locations). In the simulations of tasks from (Mushiake et al., 2006) this preference was used to setup target goal states.
- 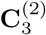: preferences over execution order cue 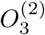 (e.g., ‘Forward’ vs. ‘Backward’).

#### Prior beliefs

The tensor 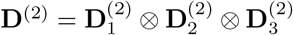 encodes priors over hidden states:

- 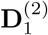: prior over *Controls* (initial policy or actions distribution).
- 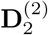: prior over *Locations* (starting position).
- 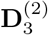: prior over *Context* (execution order).

Additionally, **E**^(2)^ encodes the prior distribution over policies. In line with the simulations from (Chen et al., 2024), where sequences vary in length, we assign equal prior probability to policies of length 1, 2, and 3. This compensates for the combinatorial bias favoring longer sequences (which are more numerous), ensuring that, for instance, upon presentation of the first target, there remains an equal prior likelihood that the full sequence will be of length 1, 2, or 3. Tab 1 shows a summary of main settings for the tasks simulated in the paper.

Note that, for simplicity, the generative models used in this study were predefined, reflecting the fact that monkeys had extensively practiced the tasks before neural recordings. However, in principle, such models could also be learned through interaction with the environment, as demonstrated in previous work (Friston et al., 2016, 2024; Van de Maele et al., 2024; George et al., 2021).

### 4.4 Worked examples of the inferential process

A schematic illustration of the inferential planning process in Section 2.2 is as follows. The start location (green; *S*_33_) is first transiently processed by early afferent neuronal populations representing locations at Level 1, whose activation is projected through the feedforward pathway into the Level 2 neuronal population. Both Level 1 and Level 2 neurons do not simply read out stimuli; instead, they implement probabilistic inference of the most probable latent causes of the sensory input by treating incoming sensory-driven activity as samples or sufficient statistics of the latent variable distribution. The feedforward pathway corresponds to the likelihood mapping (**A**) between stimuli and Level 1 hidden states, and between Level 1 and Level 2 hidden states. Lateral connections at Level 2 encode priors and transition dynamics (**B**) across hidden states over time (e.g., the effects of performing a certain action in a given state). Hidden state inference at Level 2 converges to a posterior belief via local message passing, through the continuous interaction of bottom-up evidence and internally maintained priors (see below).

Policy inference at Level 2 operates over the inferred latent states, selecting action sequences that fulfill the task goal—here, reaching the goal location (red; *S*_21_)—which is encoded as a prior preference over future observations (**C**). This is implemented by scoring candidate policies according to their (negative) expected free energy (**G**), which quantifies the divergence between predicted and preferred observations. Normally, in active inference, the first action of the selected policy would be executed, a new sensory observation gathered, and the inferential planning process repeated iteratively until the goal location is reached. However, to mimic the setup of (Mushiake et al., 2006, Figure 3), in our simulation we restrict the simulation to the planning phase. Note that in this implementation of planning dynamics, there is not a spread of activity backwards from the goal position in the sense of backwards induction [although there are active inference approaches in which this is done—e.g., (Kaplan & Friston, 2018; Friston et al., 2025)]. Instead, one starts from a selection of plausible paths (i.e., policies), and evaluates the anticipated evolution of the hidden states along each path, evaluating which of these paths is associated with the lowest expected free energy to determine its plausibility. This plausibility here is largely dominated by reaching the goal state. This means the goal state effectively sets the plausibility of the path itself, and thereby influences the probabilities of hidden states on a path to the goal via the averaging of hidden state probabilities under the policy probabilities. This has the appearance of a backwards influence, despite this never being backpropagated directly in state-space.

The simulations of forward sequences in Sections 2.3 and 2.4 follow the same scheme. The key difference is that the observed sequence itself is specified as the goal (i.e., as a prior preference over observations). For example, if the observed target sequence is *A, B, C*, these observations are encoded as preferred outcomes, which in turn drives inference toward a Level 2 (forward) policy corresponding to the action sequence *ABC*. Conversely, for backward sequences, the reversed observed sequence is specified as the goal, driving inference toward a Level 2 (backward) policy.

As discussed in Section 4.2, the inferential process is realized through message passing. In active inference, message passing is implemented through recurrent neural dynamics that continuously minimise variational free energy via local computations as set out explicitly through Equation 4.2. The local computations imply that each neural population receives messages only from neighbouring nodes in the generative model shown in Figure 5. Each neural population encodes a conditional sufficient statistic (e.g., expected hidden states or causes) and updates its activity by integrating information from other nodes, reflecting bottom-up sensory evidence or top-down predictions. These interactions give rise to recurrent error-correction dynamics, in the sense that the gradients of free energy with respect to each sufficient statistic can be taken as ‘error’ neurons that drive changes in the membrane potentials of the neurons representing those statistics. The resulting dynamics implement approximate Bayesian inference through distributed, coupled neural updates that converge toward a stationary point corresponding to the posterior belief over hidden states and policies.

The ‘message passing’ that results from this formulation is due to a ubiquitous property of the generative models used here (and elsewhere). This is a sparsity structure in which some pairs of random variables have a directional conditional dependence on one another, while others are conditionally independent. This has an important consequence for the structure of the free energy functional. If one decomposes these probabilities into a product of factors, in which each factor includes only the variables that are conditionally dependent upon one another (technically, those contained in one another’s Markov blankets (Pearl, 1988)), then this implies a sum of such terms in the free energy (due to its logarithmic representation). This means the errors associated with a given population, depending upon the gradients of the free energy, will be determined by predictions inheriting from (representations of) the variables in that population’s Markov blanket only [this is the same principle that underlies sparse message passing schemes in machine learning, including variational message passing (Winn & Bishop, 2005) and belief propagation (Yedidia et al., 2005)]. In other words, the sparsity of a generative model implies that dynamics of the sort set out in Section 4.2 are determined, for each population, by ‘predictions’ derived from a subset of other populations—giving the appearance of distributed synaptic message passing between these populations (Parr & Friston, 2018a; Parr et al., 2019b; Schwöbel et al., 2018).

### 4.5 Subspace analysis

The procedures for generating the 2D plots follow the approach described in (Xie et al., 2022). Neural responses were obtained by performing a linear regression of spike counts during the planning period against one-hot encoded task vectors (e.g., an 18-dimensional vector representing 6 directions across 3 steps). This yielded, for each neuron, a set of regression coefficients *β*(*r, l*), where *r* denotes the step and *l* the direction. For clarity, we focus on three time steps and six directions, although the procedure generalizes to other configurations (e.g., 6 directions and 2 steps, see Fig. 5).

To capture variance in neural responses—attributable to item variation at each step—we applied principal component analysis (PCA) to the regression coefficients grouped by step. This analysis produced a reliable state-space representation that captures both the relationships among step-specific subspaces and the geometry of spatial representations within each subspace.

A multivariable linear regression model was used to determine how task variables influence the average neural response during the late delay period (1 s before the go signal). For example, a length-3 sequence can be represented as an 18-dimensional three-hot vector. A sequence of targets ECA corresponds to indices [5, 3, 1] and can be encoded as:

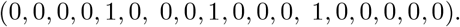

We defined 18 one-hot vectors *S*_*r,l*_ as task variables, where *r* ∈ {1, 2, 3} and *l* ∈ {1, …, 6 }. The average neural response of the *i*^th^ neuron in one trial during the late delay was modeled as:

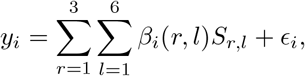

where *β*_*i*_(*r, l*) are regression coefficients and *ϵ*_*i*_ is trial-by-trial noise. To prevent overfitting, we applied Lasso regularization and selected the regularization amplitude via maximum likelihood.

The regression coefficients ***β***(*r, l*) were then used to identify low-dimensional subspaces capturing most task-related variance. Specifically, with *N* neurons, an *N* -dimensional vector

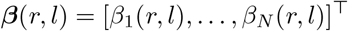

represents each rank–item combination (*r, l*) at the population level. To capture variance due to item differences at each rank, the 18 vectors ***β***(*r, l*) (*r* = 1, 2, 3; *l* = 1, …, 6) were divided into three groups by rank. For each group (fixed *r*), PCA was performed to extract the first two principal components, providing a compact representation of spatial geometry within each step-specific subspace.

### 4.6 Simulating firing rates and cosine tuning

This section explains how the firing rates shown in the simulated raster plots (e.g., Figure 2E,H) are obtained. At each update step of active inference, the model maintains: (i) a posterior over policies, *Q*_*t*_(*π*), and (ii) policy-conditioned posteriors over hidden states for each hidden-state factor *f*, denoted 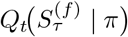, where *τ* indexes the represented time point within the plan. In the neuronal interpretation of active inference (Friston et al., 2017a), these policy-conditioned posteriors correspond to the average activity of neural populations encoding the possible hidden states under each candidate policy.

The population quantity *x*_*n*_ is obtained by Bayesian model averaging over policies:

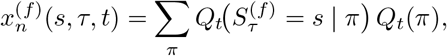

where *s* indexes the possible states of factor *f, τ* is the represented time point within the sequence, and *t* is the current update step. Thus, *x*_*n*_ is not the raw policy-conditioned posterior itself; rather, it is the policy-averaged neural readout of those beliefs. Intuitively, it expresses how strongly a population coding state *s* at sequence position *τ* is supported after averaging over all candidate policies weighted by their current posterior probability.

The simulated firing rates shown in the figures were obtained by reading out these posterior beliefs over hidden control states during inference. For each trial, the time-varying activity associated with the relevant hidden-state factor was extracted at every variational update and concatenated across task epochs, yielding one activity trace for each control-state/sequence-position combination.

For the simulations of saccadic sequences ((Figures 3, 4, 5)), only the populations corresponding to the six targets were retained. These readouts were then temporally smoothed around the stimulus epochs and transformed into target-selective population responses with a circular cosine-tuning rule applied independently within each time step (or rank). This method was used for simulating saccading sequences, but not for simulating cursor movements (Figure 2), where the cursor-movement readout was instead based directly on the policy-weighted posterior trajectories of the movement populations across sequence positions.

This tuning is obtained by assuming *B* evenly spaced preferred directions on the circle, with angular step

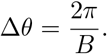

If *θ*_pref_ is the preferred direction of a given population and *θ*_*j*_ is the direction associated with the *j*-th target, the tuned response can be written as

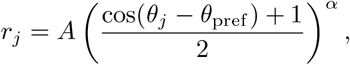

where *A* sets the peak amplitude and *α* controls the sharpness of tuning. In the six-target case used in the saccadic simulations, *B* = 6 and therefore ∆*θ* = *π/*3 (i.e., 60^*◦*^). In the simulations, we used a peak amplitude of *A* = 0.85 and a sharpness parameter *α* = 5, yielding a peak at the preferred direction and a marked decay over neighbouring directions.

To generate the firing-rate panels, each population trace was expanded into a small ensemble of nominally identical neurons, so that multiple neurons shared the same mean response profile for a given control state and time step. The raster plots were then obtained by converting these continuous firing rates into binary spike trains on a fine temporal grid. A short refractory silence was imposed after each emitted spike, preserving the slower inferential dynamics of the underlying firing-rate trajectories.

The spike generator used for the sequence simulations treated the firing rate itself as a spike probability. If *F*_*i,t*_ ∈ [0, 1] denotes the firing rate of neuron *i* at time bin *t*, spikes are sampled as independent Bernoulli events:

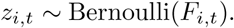

Finally, a refractory rule is applied: if *z*_*i,t*_ = 1, then the following *r* bins are forced to zero,

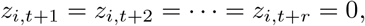

thereby preventing unrealistically dense spike bursts in adjacent bins.

In the current extraction settings, the refractory parameter was *r* = 3 bins.

### 4.7 Dimensionality estimation and participation ratio

To summarize the dimensionality of population activity, we used the participation ratio, often interpreted in the literature as an effective dimensionality measure (Gao et al., 2017; Recanatesi et al., 2022). The participation ratio, in fact, provides a global estimate of effective dimensionality based on the eigenspectrum of the covariance matrix. If *λ*_*i*_ are the eigenvalues of the covariance matrix of the population activity, the participation ratio is

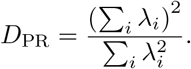

This quantity is large when variance is distributed across many components and smaller when variance is concentrated in a few dominant components.

## Acknowledgments

This research received funding from the European Research Council under the Grant Agreement No. 820213 (ThinkAhead), the Italian National Recovery and Resilience Plan (NRRP), M4C2, funded by the European Union – NextGenerationEU (Project IR0000011, CUP B51E22000150006, ‘EBRAINS-Italy’; Project PE0000013, ‘FAIR’; Project PE0000006, ‘MNESYS’), and the Ministry of University and Research, PRIN PNRR P20224FESY and PRIN 20229Z7M8N. The GEFORCE Quadro RTX6000 and Titan GPU cards used for this research were donated by the NVIDIA Corporation. T.P. is supported by an NIHR Academic Clinical Fellowship [ref: ACF-2023-13-013]. KF is supported by funding from the Wellcome Trust (Ref: 226793/Z/22/Z). J.C.R.W is supported by European Research Council Starting Grant No. 101222868 (NARFB). We used a Generative AI model to correct typographical errors and edit language for clarity.

## Notes

### Competing Interest Statement

The authors have declared no competing interest.

### Summary of Updates

Figures 2 to 5 revised, text corrected, supplementary materials included

