## Supplementary Materials for "Inferential planning in the frontal cortex"

### Supplementary Material for
### Inferential planning in the frontal cortex

Francesco Donnarumma[1] Thomas Parr[2] Karl Friston[3,4] James Whittington[5]
Giovanni Pezzulo[1,*]

[1] Institute of Cognitive Sciences and Technologies, National Research Council, Rome, Italy
[2] Nuffield Department of Clinical Neurosciences, University of Oxford, Oxford OX1 2JD, UK
[3] Queen Square Institute of Neurology, University College London, London WC1E 6BT, UK
[4] VERSES Research Lab, Los Angeles, CA 90016, USA
[5] Department of Experimental Psychology, University of Oxford, Oxford, UK

June 5, 2026

### Policies used in the four simulations

In the simulations presented in the main text, a policy at Level 2 is defined as a sequence of control states. Table 1 summarizes the policies considered in each simulation. Additional details for each simulation are provided in the sections below.

| Simulation | Policies | Number of policies |
|---|---|---|
| Simulation (1):<br>Cursor movements<br>(Mushiake et al., 2006) | All the policies have length $T^{(2)} = 3$<br>The start state is always $S_{33}$<br>The goal state varies from trial to trial | 16 |
| Simulation (2):<br>Sequence of saccades<br>(Xie et al., 2022) | All the policies have length $T^{(2)} = 3$<br>Targets are $\{A, \ldots, F\}$, without repetition | $6P3 = 120$ |
| Simulation (3):<br>Variable-length forward/<br>backward sequence of saccades<br>(Chen et al., 2024) | Policies have lengths $1, 2, 3$<br>Targets are $\{A, \ldots, F\}$, without repetition<br>Two subsets of policies for *Forward* and*Backward* sequences<br>Context (forward or backward) known in advance | $6P1 + 6P2 + 6P3 = 156$ forward<br>$6P1 + 6P2 + 6P3 = 156$ backward<br>Total $= 312$ policies |
| Simulation (4):<br>Fixed-length forward/<br>backward sequence of saccades<br>(Chen et al., 2024) | All the policies have length $T^{(2)} = 2$<br>Targets are $\{A, \ldots, F\}$, without repetition<br>Two subsets of policies for *Forward* and *Backward* sequences<br>Context (forward or backward) inferred based on a cue $t_{cue}$ | $6P2 = 30$ forward<br>$6P2 = 30$ backward<br>Total $= 60$ policies |

**Table 1 Policies used in the four simulations.** See the subsequent sections for additional details.

In the table, $nPk$ denotes the number of dispositions/permutations of $k$ elements chosen from $n$, with order relevant:

$$nPk = \frac{n!}{(n-k)!}.$$

For example, 6P3 in Simulation (2) denotes the number of permutations of length 3 that can be formed from 6 targets. Since all policies have length 3 and targets cannot be repeated, this yields a total of 120 possible policies.

#### Supplementary information for Simulation (1)

Simulation (1) addresses the task of (Mushiake et al., 2006), requiring monkeys to plan a sequence of cursor movements from the central start state $S_{33}$ to a known goal state that changes for each problem. Note that only some policies are allowed, given the configuration of the walls. This information is encoded as a policy prior, $\mathbf{E}^{(2)} = p(\pi \mid \text{Maze})$. In our simulations, the policies comprise all valid three-step paths under the maze topology, excluding paths that revisit the same state. This yields 16 admissible policies:

`URR, ULL, DRR, DLL, URD, ULD, DRU, DLU,`
`UUR, UUL, DDR, DDL, DRD, DLD, URU, ULU.`

For each problem, the goal state is set as a preferred state (i.e., as a very high prior over state-specific observation) and the best policy is inferred using standard active inference methods (Parr et al., 2022).

Supplementary Figure S1 shows an illustrative simulation of the planning process depicted in Figure 2G–I of the main text, in which the policy $U(1), R(2), R(3)$ is inferred from the start location $S_{33}$ to the goal location $S_{25}$, complementing the results shown in the main text. It illustrates firing rates and simulated raster plots associated with hidden state inference at Level 2 (Figure S1A,C) and Level 1 (Figure S1B,D). The Level 2 inference is identical to that shown in the main text, but here only the firing rates associated with the inferred plan are displayed. The Level 1 inference illustrates the dynamics of hidden states associated with actions to be executed. The figure highlights the distinction between persistent and parallel activation of multiple elements of an action plan at Level 2 and the transient, sequential activation of inferred actions at Level 1. Note that the actions inferred at Level 1 during the planning process could be executed after planning. Indeed, in standard active inference simulations, planning (inference of a policy) is interleaved with action execution (typically executing the first action of the inferred policy), followed by the acquisition of a new observation and subsequent replanning, until the goal is achieved (Parr et al., 2022). The execution phase is not shown here but would follow the same transient and sequential dynamics illustrated at Level 1 (Figure S1B,D). This is because inferred actions are executed sequentially, one at a time, thereby producing transient, sequential dynamics at Level 1.

Supplementary Figure S2 shows the dynamics of another component of the generative model not shown in the main text—namely, the hidden location states—during inference of the policy $U(1), R(2), R(3)$, at both Level 2 (Figure S2A,C) and Level 1 (Figure S2B,D), using the same format as Figure S1. The hidden location states correspond to the 21 possible locations of the maze, labeled from $S_{12}$ to $S_{54}$ as in Figure 2G of the main text. The figure highlights the distinction between persistent and parallel activation of multiple locations—corresponding to the locations expected to be visited while executing the inferred policy—at Level 2 and the transient, sequential activation of hidden states encoding inferred locations at Level 1. Note that the former are not mandatory but

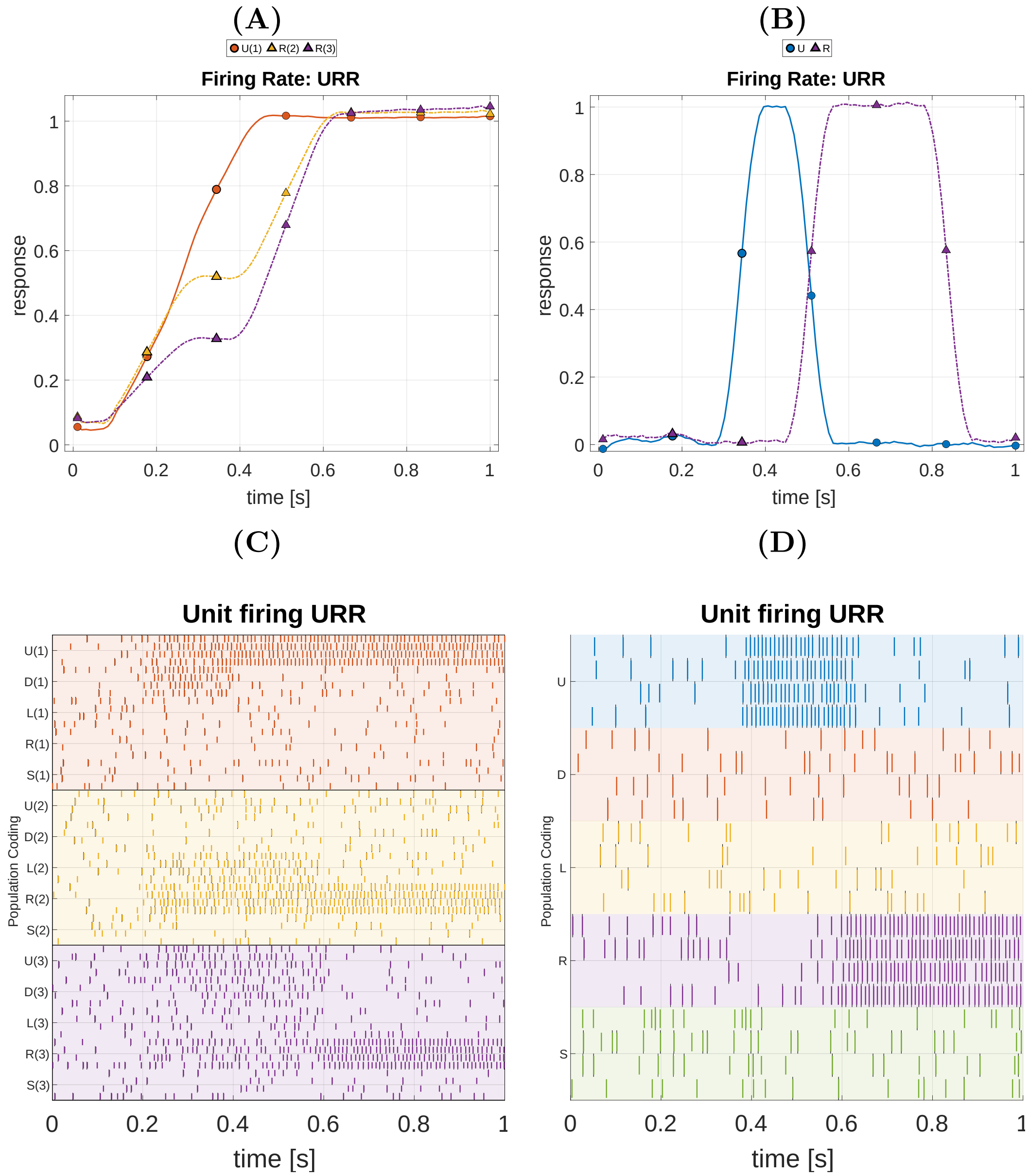


**Figure S1 Extended Simulation (1): hidden control states at both levels of the hierarchical generative model during inference of the $U(1), R(2), R(3)$ policy.** This figure shows the neural readouts associated with hidden control states (actions) during the planning phase. Panels (A) and (B) report firing rates associated with hidden state inference at Level 2 and Level 1, respectively. Panels (C) and (D) show the corresponding raster plots at Level 2 and Level 1. See the main text for further explanation.

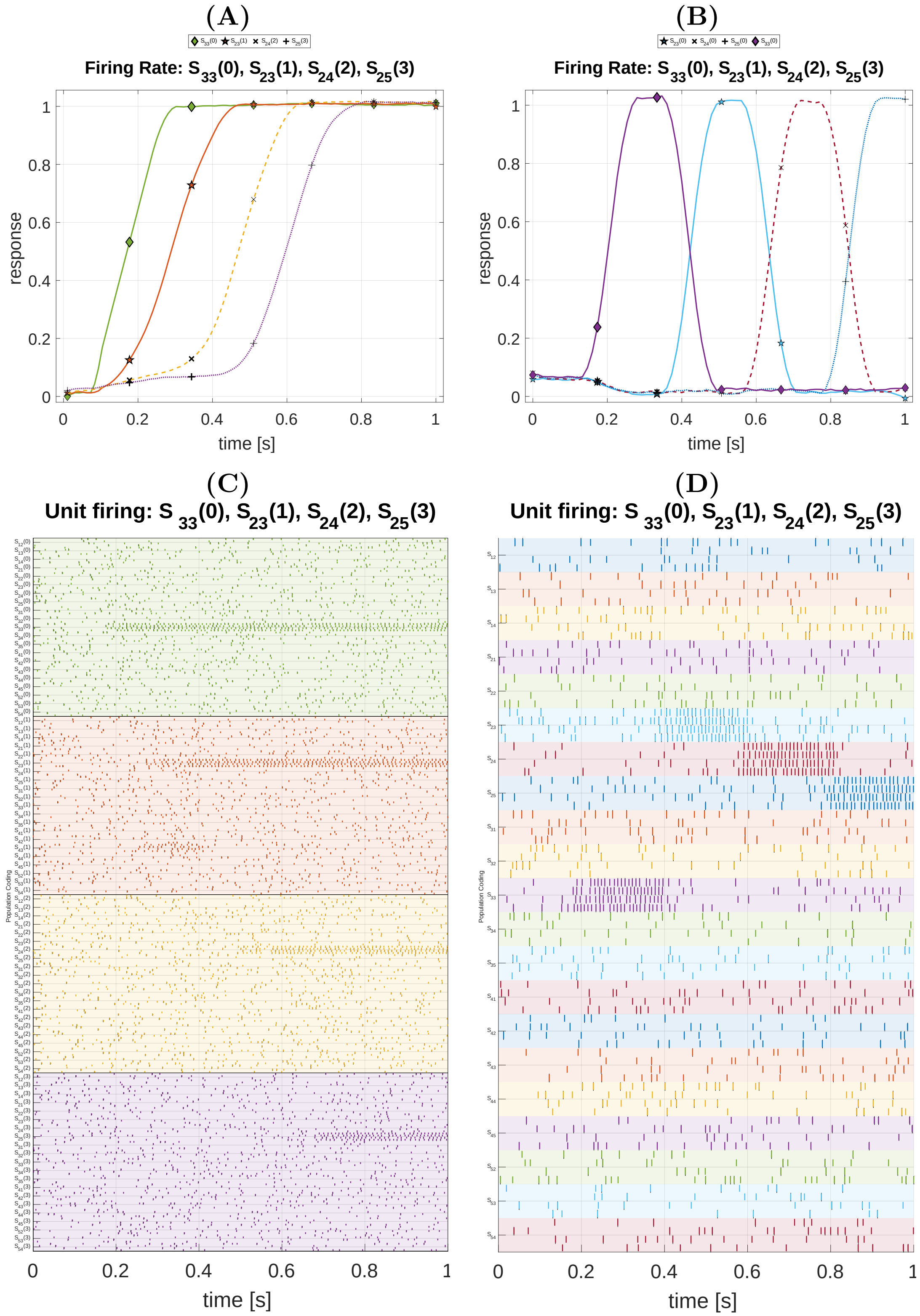


**Figure S2 Extended Simulation (1): hidden location states at both levels of the hierarchical generative model during inference of the $U(1), R(2), R(3)$ policy.** This figure shows the neural readouts associated with hidden location states during the planning phase. Panels (A) and (B) report firing rates at Level 2 and Level 1, respectively. Panels (C) and (D) show the corresponding raster plots at Level 2 and Level 1. See the main text for further explanation.

#### Supplementary information for Simulation (2)

Simulation (2) addresses the task of (Xie et al., 2022), in which monkeys are required to infer and maintain in memory a sequence of three saccades.

In this simulation, policies correspond to ordered sequences of three saccadic targets drawn from the six possible targets $A$ to $F$, without repetition. Examples include $ABC$, $ABE$, and $ABF$. The number of distinct policies is therefore $6P3 = 120$.

#### Supplementary information for Simulation (3)

Simulation (3) addresses the main task of (Chen et al., 2024), in which monkeys are required to infer and maintain in memory a variable-length sequence of saccades, which they later execute in either forward or backward order. Note that the monkeys are explicitly instructed whether the execution context is forward or backward.

In this simulation, policies correspond to variable-length target sequences of lengths 1, 2, and 3. For each execution context, the number of sequences is

$$6P1 + 6P2 + 6P3 = 6 + 30 + 120 = 156.$$

The full policy space therefore consists of two sets of 156 policies, one for the *Forward* context and one for the *Backward* context, for a total of 312 policy–context combinations.

Because the execution context is already known at the beginning of the task, we set a prior over policies that selects from the outset the relevant subset (*Forward* or *Backward*) rather than mixing both during inference. As stated in the main Methods, the prior over policies assigns equal total probability mass to policies of lengths 1, 2, and 3, compensating for the larger number of longer sequences so that the three length classes are equiprobable a priori.

Supplementary Figure S3 shows representative examples of the forward and backward contexts when the sequence length is two (i.e., the last target is omitted). This simulation complements the examples provided in the main text, where the sequence length is three.

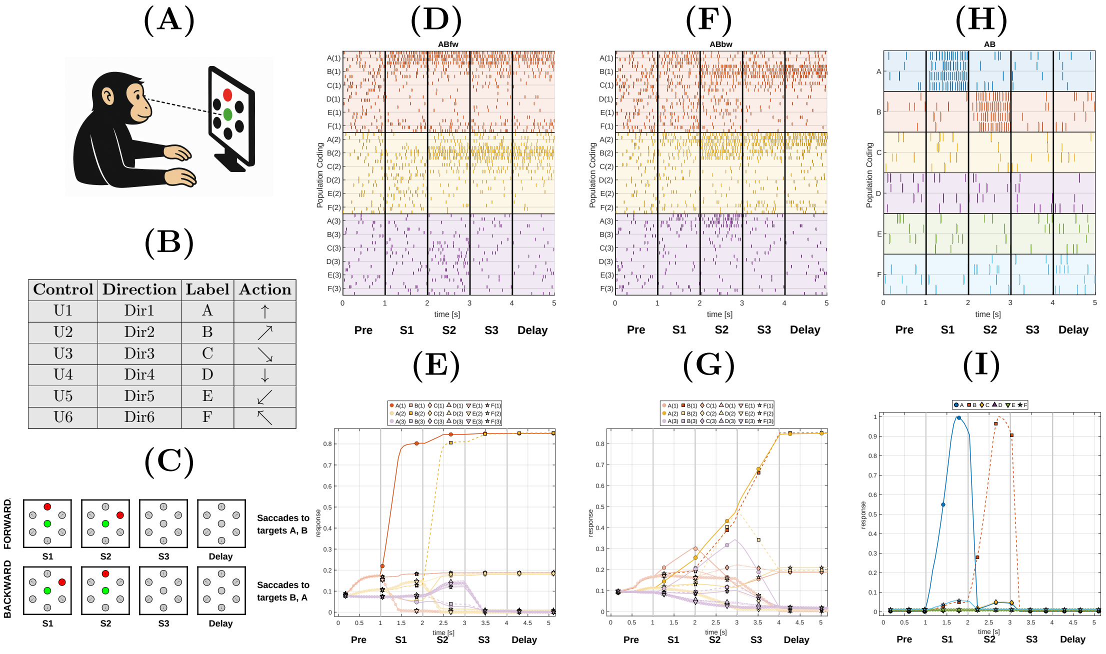


**Figure S3 Extended Simulation (3): inferring forward and backward sequences of two targets.** This figure refers to the variable-length saccade experiment of (Chen et al., 2024). Panels (A) and (B) show the saccade task and the mapping between control states, target labels, and movement directions. Panel (C) illustrates the case of a sequence of length two: targets are presented at S1 and S2, S3 is omitted, and the inferred sequence is maintained during the delay period. Panels (D) and (E) show a Forward-context simulation for a two-target sequence. (D) Raster plot of simulated neural activity, with four neurons coding each of the six control states (targets A–F) at Level 2, at each of the three sequence positions. (E) The same activity plotted as simulated firing rates. Panels (F) and (G) show the corresponding Backward-context simulation in the same format. Panels (H) and (I) show Level 1 activations during sequence observation, again in raster and firing-rate format. See the main text for further explanation.

#### Supplementary information for Simulation (4)

Simulation (4) addresses a task of (Chen et al., 2024), in which monkeys are required to infer and maintain in memory a fixed-length sequence of two saccades, which they later execute in either forward or backward order. Note that the monkeys are not instructed whether the execution context is forward or backward, but instead must infer it based on a cue observed after the targets.

In this simulation, policies correspond to ordered target sequences of length 2, again without repetition, giving $6P2 = 30$ base sequences. However, the same pair of targets can be executed either in the *Forward* or *Backward* direction, depending on the cue observed at $t_{\mathrm{cue}}$. For this reason, the full space comprises 60 policy–context combinations. In this simulation, the policy prior is also asymmetric before the cue: *Forward* policies are made one order of magnitude more probable than *Backward* policies ($w_{\mathrm{Forw}} = 10\, w_{\mathrm{Back}}$). Consequently, before $t_{\mathrm{cue}}$ the posterior tends to be dominated by *Forward* policies; after the cue, inference uses the new sensory evidence to correct this initial bias when *Backward* execution is required.

To illustrate the role of the strong prior for *Forward* policies, Supplementary Fig. S4 shows the same task in a symmetric variant in which *Forward* and *Backward* policies have equal prior weight. In this case, the model exhibits a correspondingly symmetric inferential behaviour across the two execution contexts, which is not observed empirically in (Chen et al., 2024).

It might be argued that the coexistence of two different targets within the same "memory subspace" would lead to information interference, if a neural population were attempting to decode the content of the subspace. However, in our framework, the plan is inferred, not decoded, and the coexistence of two targets is simply a signature of inference under equal prior probability. It therefore does not impair interference, as evidenced by the model's ability to recover the correct plan once the forward or backward cue is observed.

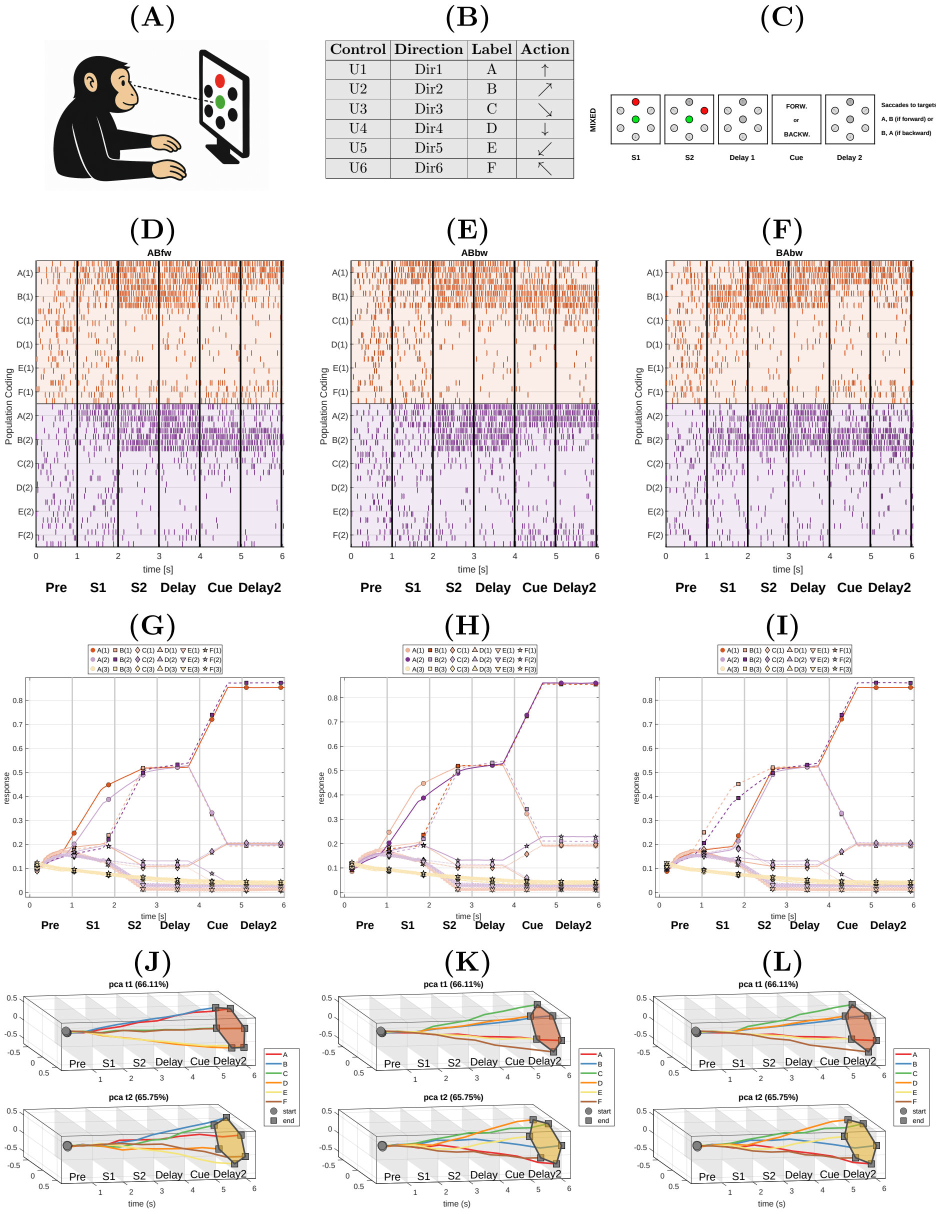


| Control | Direction | Label | Action |
|---|---|---|---|
| U1 | Dir1 | A | ↑ |
| U2 | Dir2 | B | ↗ |
| U3 | Dir3 | C | ↘ |
| U4 | Dir4 | D | ↓ |
| U5 | Dir5 | E | ↙ |
| U6 | Dir6 | F | ↖ |

**Figure S4 Extended Simulation (4). Exploring the role of an equal prior for *Forward* and *Backward* policies.** This supplementary figure follows the same task structure as the fixed-length saccade experiment of (Chen et al., 2024), but here *Forward* and *Backward* policies are given equal prior weight. Panels (A)–(C) summarize the task structure corresponding to the explanatory panels of Fig. 5 in the main text: two targets are presented sequentially at S1 and S2, followed by a delay period and an instruction cue indicating whether the sequence should be executed in the forward or backward direction. Panels (D)–(F) show raster plots for three example cases: AB in the forward condition, AB in the backward condition, and BA in the backward condition. Panels (G)–(I) show the corresponding simulated firing rates. Panels (J)–(L) show the corresponding trajectories of the first two principal components. In contrast to Figure 5 in the main text, the absence of a strong *Forward* prior yields a more symmetric behaviour across forward and backward conditions, which is not observed empirically in (Chen et al., 2024). See the main text for further explanation.
